# Spatial chromosome organization and adaptation of the radiation-resistant extremophile *Deinococcus radiodurans*

**DOI:** 10.1101/2023.11.19.567671

**Authors:** Qin-Tian Qiu, Cai-Yun Zhang, Zhi-Peng Gao, Bin-Guang Ma

## Abstract

Radiation-resistant *Deinococcus radiodurans* is an extremophilic microorganism capable of withstanding high levels of ionizing radiation and chemical mutagens. It possesses remarkable DNA repair capability and serves as a model organism for studying stress resistance mechanism. However, our understanding on the spatial chromosome organization of this species remains limited. In this study, we employed chromosome conformation capture (3C) technology to determine the 3D genome structure of *D. radiodurans* and to further investigate the changes of chromosome conformation induced by ultraviolet (UV) irradiation. We observed that UV irradiation reduced short-range chromosome interactions, and smaller chromosomal interaction domains (CIDs) merged to form larger CIDs. Integrating transcriptomic data analysis, we found that the majority of upregulated differentially expressed genes were significantly enriched near specific CID boundaries. Specially, we comprehensively elucidated that the nucleoid-associated protein Dr_ebfC may serve as a global regulator for gene expression by altering chromosome structure, thereby influencing the physiological state of the bacterium. Overall, our study revealed the chromosome conformations of *D. radiodurans* under different conditions, and offered valuable insights into the molecular responses of this extremophile to environmental stresses.

## Introduction

For both eukaryotes and prokaryotes, the carrier of genetic information, DNA, combines with proteins to form chromosome which requires a high degree of compression to fit into a cellular space thousands of times smaller than the length of stretched DNA molecule. The highly compacted chromosomes form organized structures to meet the functional requirements of gene expression, DNA replication, and chromosome segregation (1). Over the past decade, significant advancements in our understanding of bacterial genome dynamics and interactions have been achieved through the application of chromosome conformation capture (3C)-based techniques (2–9). Presently, chromosome contact maps of each studied bacterial genome have shown clear chromosomal interaction domains (CIDs), similar to the topologically associating domains (TADs) found in eukaryotes (10,11). Chromosome regions within the same CID tend to interact more frequently than inter-domain regions, and directionality index (DI) or insulation score can be used to identify such domains based on this characteristic. Except for the smaller-sized domains (15-30 kb) found in *Mycoplasma pneumoniae*, CIDs in other bacteria typically range from 30 to 400 kb (5). The specific mechanisms underlying CID formation are still unclear. It is suggested that the formation of CID may be related to transcription process, DNA supercoiling, and macromolecular crowding. It has been observed that CID boundaries almost disappear after treatment with the transcription inhibitor Rifampicin in *Bacillus subtilis*, suggesting that long and active transcriptional regions contribute to CID boundary formation (2,12). In addition, nucleoid-associated proteins (NAPs) play diverse roles in the process of bacterial nucleoid organization (13,14). In *Escherichia coli*, various NAPs act on chromosomal structures at local or global scales. For example, HU and Fis proteins facilitate long-range DNA interactions, while the MatP protein restricts chromosomal interactions within a 280 kb region at the chromosome replication terminus region (6). A recent study found that Rok protein in *B*. *subtilis* forms large (Mb range) and well-defined anchored chromosomal loops, physically isolating large regions of chromosome (15).

*D. radiodurans* exhibits an exceptional capacity to withstand extreme environmental conditions, characterized by a radiation tolerance approximately 30-fold greater than that of *E. coli* and several thousand-fold greater than that of human cells (16). Generally, radiation can disrupt the structure and function of chromosomal DNA through physical effects and secondary oxidative damage, resulting in a substantial accumulation of single or double stranded DNA breaks, which culminate in cellular demise. *D. radiodurans* possesses 4-10 genome copies within a single cell, and studies have suggested that its robust environmental adaptation capabilities are largely attributed to its multiple genome copies or redundant genetic information (17). Despite extensive researches on DNA damage repair mechanism under environmental stress (18–22), the investigation of chromosomal 3D structures in response to stress in *D. radiodurans* is still lacking. Furthermore, compared to radiation-sensitive bacteria, *D. radiodurans* possesses highly compacted nucleoid structures that remain unchanged even after exposure to high doses of radiation (23). NAPs may play a role in this high degree of nucleoid compaction. The types and quantities of NAPs in *D. radiodurans* are species-specific (24). Based on amino acid sequence analysis, a few known NAPs have homologs in *D. radiodurans*, including HU (*dr_a0065*), DnaA (*dr_0002*), Dps (with *dr_2263* referred to as Dps1 and *dr_b0092* as Dps2), Lrp (*dr_1894*, *dr_0200*), and EbfC (*dr_0199*) (25–28).

EbfC (*erp*-binding factor, chromosomal), also known as YbaB, is a less-studied NAP but widely distributed in prokaryotes (29). It was first discovered in *Borrelia burgdorferi* and characterized as a site-specific DNA binding protein (30). Homologs of EbfC found in *E. coli* and *Haemophilus influenzae* also function as a transcriptional regulator and play roles in DNA repair. The EbfC protein in *D. radiodurans*, identified in 2012, is a novel member of the NAP family. Preliminary experiments have shown that the Dr_ebfC protein localized in the nucleoid region of cells and constrained DNA supercoiling *in vitro* (28). So far, research on the YbaB/EbfC family proteins has mainly focused on exploring their functions using molecular biological methods. From the perspective of 3D genome, further investigations are required to unravel the role of EbfC in the mechanism of chromosome organization.

In this study, the chromosome conformation capture and sequencing (3C-seq) technique was applied to explore the structural features of *D. radiodurans* chromosomes, and the differences in chromosome 3D structure and gene expression before and after ultraviolet (UV) irradiation were analyzed combined with transcriptome data. Furthermore, the impact of Dr_ebfC protein inactivation on chromosomal contacts was elucidated, manifested by the reduction in local short-range interactions and changes in gene expression level resulting from chromosome structural alterations. Findings in this research for the first time reveal the effects of UV irradiation and the NAP Dr_ebfC on chromosome organization at 3D genomics level, and provide a new perspective for understanding the role of chromosome organization in the radiation resistance mechanism of *D. radiodurans*.

## Materials and methods

### Strain, growth conditions and treatment

All strains, plasmid used in this study are listed in **Table S1**. *D. radiodurans* strains were grown at 30°C in tryptone glucose yeast extract (TGY) liquid media or on agar plates (0.5% tryptone, 0.1% glucose, and 0.3% yeast extract). The cultures were grown to OD_600_ = 3.0 (middle exponential). For UV irradiation, the experiment was performed as reported previously (31).

### Construction of the *D. radiodurans* ΔDr_ebfC strain

The ΔDr_ebfC (*dr_0199*) strain was constructed by a tripartite ligation method, as described previously (32). Briefly, the DNA fragment upstream of *dr_0199* was amplified by PCR using the primers Δ*dr_0199*-p1 and Δ*dr_0199*-p2, which was digested with BamHI (**Table S1**). The DNA fragment downstream of *dr_0199* was amplified by PCR using the primers Δ*dr_0199*-p3 and Δ*dr_0199*-p4, which were digested with HindIII (**Table S1**). The digested fragments were ligated to a kanamycin resistance gene that was digested with BamHI and HindIII previously (**Table S1**). After the triplet ligation product was transformed into the *D. radiodurans* wild-type R1 strain, the mutant colonies were then selected on TGY plates containing 20 µg/ml kanamycin, and confirmed by genomic PCR using two pairs of primers (Δ*dr_0199*-p1 and Δ*dr_0199*-p4, Δ*dr_0199_test-*F and Δ*dr_0199_test-*R) and DNA sequencing.

### Growth curves and phenotypic assay

To measure the growth curve, the wild-type *D. radiodurans* R1 and ΔDr_ebfC strains were pre-cultured to OD_600_ = 3.0 at 30°C with shaking at 220 rpm/min and then 1 ml of culture was transferred into 100 ml of fresh TGY medium without antibiotics. OD_600_ values were monitored at 2-hour intervals until a point was reached where there was no longer a significant increase in the measured values. The final growth curve corresponding to the mean of three independent experiments was plotted using Origin 2023 software. To observe phenotypes under H_2_O_2_ treatment, the wild-type *D. radiodurans* R1 and ΔDr_ebfC strains were pre-cultured to OD_600_ = 3.0, then treated with 50 mM H_2_O_2_ for 30 min. After the reaction, the mixture was diluted in gradient and plated on TGY plate. The experiment was repeated three times.

### Transcriptome data analysis

Bacterial cells were pre-cultured in TGY to an OD_600_ = 3.0, harvested by centrifugation at 12,000× g for 3 min and stored at −80°C. Cryopreserved cells were dispatched to Wuhan Frasergen Bioinformatics Co., Ltd., where RNA extraction, cDNA library construction, and on-machine sequencing procedures were carried out. The sequencing standard was paired-end 150 bp, and the sequencing platform was MGISEQ-2000. After removing the adapter sequences and low-quality sequences from the raw reads, clean reads were obtained for mapping to the *D. radiodurans* R1 genome (GenBank assembly accession GCA_000008565.1) with Hisat2 Aligner (33). The mapped reads of each sample were applied to quantify the expression levels using StringTie (34) with a reference-based approach. Differential expression of genes was normalized and calculated by DEseq2 algorithm (35). Genes with expression level changes more than twofold increased or reduced with adjusted *p*-value < 0.05 were considered as differentially expressed genes (DEGs).

### 3C library construction and sequencing

3C experiments were carried out following the same procedure as described previously (6). Briefly, 20 mL of cell culture were collected by centrifugation and crosslinked with fresh formaldehyde for 30 minutes at 25℃ (7% final concentration) followed by 30 minutes at 4°C. Formaldehyde was then quenched with a final concentration of 0.25 M glycine for 20 minutes at 4°C. The fixed cells were collected by centrifugation and washed once with double distilled water. The cell pellets were frozen in liquid nitrogen and stored at −80°C until use.

Frozen pellets were thawed in the ice box, resuspended and lysed by incubation with lysozyme at 37°C for 30 min. The reaction was quenched by adding 10% SDS and incubating for 5 min at 65°C. 50 μL of lysed cells were then transferred in one EP tube containing 450 μL of digestion mix (355 μL of double distilled water, 50 μL of 10 × rCutSmart™ Buffer, 25 μL of 25% Triton X-100, and 20 μL of 10 U/μL HpaII enzyme). DNA was digested for 10 hours at 37°C with shaking at 200 rpm/min. For ligation, the reaction was mixed with 7945 μL of double distilled water, 1000 μL of 10 × T4 DNA Ligase Reaction Buffer (NEB), 50 μL of 20 mg/mL BSA (TaKaRa), 500 μL of 20% Triton X-100, and 5 μL of 400 U/μl T4 DNA ligase (NEB). The ligation reaction was incubated for 4 h at 16°C with occasional inversion of the tube (every 1 h). After the ligation was completed, the reaction was mixed with 100 μL of 0.5 M EDTA and 50 μL of 20 mg/ml proteinase K (TransGen) and incubated at 58°C for 4 h. Precipitation of DNA was performed by standing overnight at 4°C in presence of 3 M Na-Acetate (pH 5.2) and iso-propanol. In the next day, the DNA was extracted through using DNA Extraction Reagent (SOLARBIO). The obtained DNA was washed with 500 μL 75% cold ethanol, dissolved in 40 μl 1 × TE buffer containing 0.1 mg/ml RNase A and incubated for 30 min at 37°C. Successful proximity ligation was confirmed by running 5 μL of the DNA on a gel. Finally, the resulting DNA was used for library construction and paired-end sequenced on the BGI DNBSEQ platform. Two biological replicates were used for each condition.

### 3C-seq data analysis

3C-seq reads were filtered and quality-evaluated using software Cutadapt (36) and FastQC with the default parameters to generate clean reads. The clean reads were aligned independently to the reference genome (*D. radiodurans*: GCA_000008565.1) by Bowtie2 (37) using a very-sensitive and iterative approach. Each read was then assigned to a restriction fragment, with uninformative events, such as self-circularized and uncut fragments discarded by filtering. Because these restrictive fragments presented unequal lengths, the genome was binned into 5 kb segments to generate the interaction matrix files. After correlation analysis, the two biological replicates were merged for subsequent analysis. The original contact matrices were then normalized using the sequential component normalization procedure (SCN) (38) to reduce the biases inherent resulting from the 3C experimental protocol.

CIDs were identified by the DI according to a previous study (6). CID boundaries were defined as the area where the direction preference of bins shifted from negative values of upstream segments to positive values of downstream segments. Sequence motifs of CID boundaries were investigated in the PRODORIC database using MEME Suite (https://meme-suite.org). Significant intra-chromosome interactions in which both the *p*-value and *q*-value (adjusted *p*-value for Benjamini-Hochberg correction) were less than 0.05, and contact count > 2 were selected by FitHiC2 software (39). To identify DNA loops, we applied the Chromosight (40) “detect” function (with parameter settings --threads 8 --min-dist 20000 --max-dist 250000 --pearson 0.32) to each normalized matrix (of 5 kb bins). This algorithm output a score (Pearson coefficient) which allows for the interpretation of results and the comparison of different mutants/treatments. This value was used as the strength of the loop in this study.

### Reconstruction of the 3D chromosome model of *D. radiodurans*

The 3D chromosome models of *D. radiodurans* were reconstructed based on the SCN normalized interaction frequency (IF) matrix using the EVRC software (41). The EVRC program generates PDB files containing the spatial coordinates of each node in the 3D structure corresponding to a bin of the IF matrix. The coordinate file was visualized using PyMOL (The PyMOL Molecular Graphics System, Version 2.4.0 Schrödinger, LLC).

## Results

### Global conformation of the *D. radiodurans* genome

Chromosomal 3D structures reflect the regulatory functions of genome organization in cellular activities. In this work, we applied 3C-seq to the wild-type cells of *D. radiodurans* R1 strain in the mid-exponential phase to explore the 3D organization of its multipartite genome. The genome of *D*. *radiodurans* consists of four replicons: the large chromosome Chr1 [2649 kb], the small chromosome Chr2 [412 kb], the large plasmid pMP1 [177 kb], and the small plasmid pCP1 [46 kb] (42). **Figure 1A** displays the normalized contact map of chromosomes with a resolution of 5 kb, and it shows strong diagonal signal, indicating a propensity for neighboring loci to interact with each other along the genome. Additionally, the contact map of Chr1 exhibits a faint secondary diagonal, reflecting weak inter-arm interactions between loci on the left and right arms of Chr1, and there seems no significant inter-arm interactions for Chr2 and the two plasmids (**Fig. S1**). Compared with other bacteria, it was found that the interaction pattern of Chr1 resembles that of *B*. *subtilis*, *Caulobacter crescentus*, and *Vibrio cholerae* Chr2 (43), where the presence of secondary diagonal interactions reflects the circular nature of the bacterial genome. To further interpret the contact map, scalograms were generated by calculating the sum of interaction frequencies within a certain range upstream and downstream of each bin on the chromosome, which provide insights into local chromosomal behaviors. This visualization tool has been employed to investigate the impact of NAPs on the local spatial compaction of the genome in *E. coli* (6). **Figure 1B** illustrates the uneven distribution of local compaction of *D*. *radiodurans* Chr1 and Chr2 at different scales.

Given the small size (in bp) of the two plasmids in *D*. *radiodurans*, we focused on modeling the 3D structures of Chr1 and Chr2 at a 5 kb resolution using the EVRC software (41). This software considers both intra-chromosomal and inter-chromosomal interactions, allowing us to reconstruct this multipartite genome. **Figure 1C** shows the obvious difference in the structure of the two chromosomes. Specifically, Chr1 exhibits a boat-like shape with its replication origin (*ori*1) located at the bow of the boat in the 3D model, corresponding to the first bin (1183-1903 bp) on the genome. The arms of Chr1 align along the longitudinal axis and have a closer proximity. In contrast, Chr2 has a more circular shape, with larger distance between the arms, and its replication origin (*ori*2) is also located in the first bin of the Chr2 genome. Previous work in *E. coli* has found that protein-protein interaction (PPI) correlates with DNA interaction on a genome scale (44). To investigate this feature, we collected the PPI pairs (totally 2997) of *D*. *radiodurans* from the STRING database (45). Using the spatial distance from the chromosome 3D model, we found that DNA fragments involved in functional PPIs are spatially closer to each other (*p* = 3.6 × 10^-66^) (**Fig. 1D**). Importantly, this phenomenon was also observed when environmental condition or physiological state of this organism changed (**Fig. S2**), highlighting the conservativeness of this chromosomal 3D feature.

**Figure 1.**
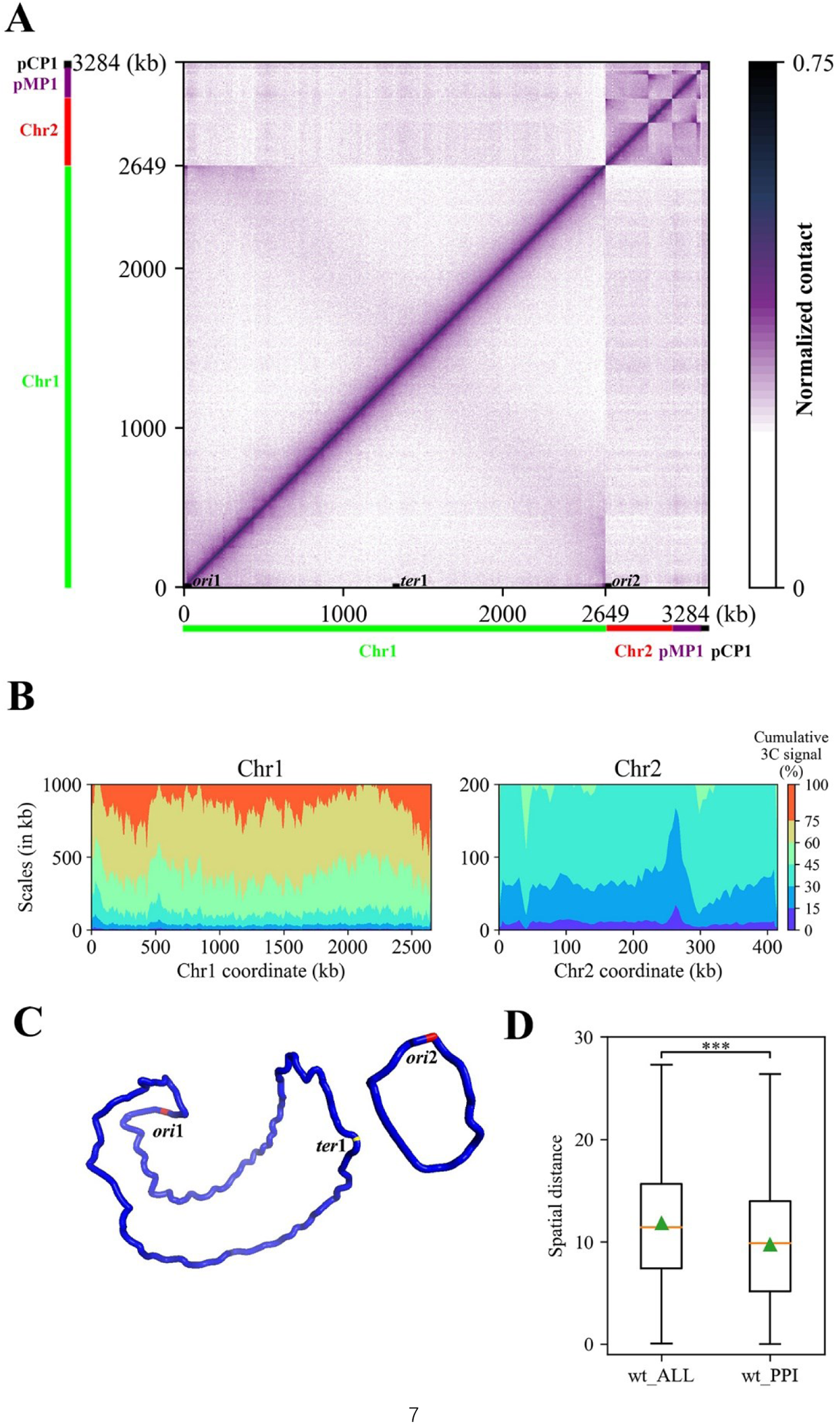
Spatial organization of *D. radiodurans* replicons. (A) Normalized contact map displays contact frequencies of 5 kb bins across the genome of *D. radiodurans* R1 wild-type (WT) cells growing exponentially. The *x* and *y* axes represent genomic coordinates of each replicon. Chr1, Chr2, pMP1, and pCP1 are indicated by green, red, purple, and black bars, respectively. The positions of the origins (ori1 and ori2) and terminus (ter1) are indicated on the *x* axis. (B) Scalogram representation of the two chromosomes of *D. radiodurans*. Scalograms reflect relative compactness of the chromosome regions. The colored areas above each bin represent the fraction of the total cumulated contacts made by the bin with its flanking regions of increasing sizes (dark blue, 0–15%; light blue, 15–30%, …; red, 75–100%). (C) The 3D model of the two chromosomes of *D. radiodurans*. The origins (*ori*1 and *ori*2) are marked as red and the terminus (*ter*1) is marked as yellow. (D) Boxplot of the spatial distance distribution between bin pairs in the 3D mode. wt_ALL: all bin pairs in the genome; wt_PPI: the bin pairs containing protein-protein interaction (namely, one bin in a bin pair contains one of the two interacting proteins, and the other bin in the bin pair contains the other interacting protein). The orange line indicates the median of the box. The green triangle indicates the average value of the box. ***: *p*-value < 0.001.

### Analysis of chromosomal interaction domains (CIDs) in the 3D genome of *D. radiodurans*

The 3C contact map exhibits highly self-interacting regions or CIDs that appear as squares along the main diagonal (**Fig. 1A**). When the map is rotated 45° clockwise, it appears as triangulars (**Fig. 2A**). Using the directionality index (DI) at a scale of 100 kb, we identified a total of 23 CIDs in the wild-type cells (Chr1: 21, Chr2: 2) (**Fig. 2B**), ranging in size from 30 kb to 250 kb (average size: 101 kb), which is consistent with the CID scales observed in other bacteria (5). Further analysis of the guanine and cytosine (G+C) content distribution within the internal and boundary regions of the CIDs in *D*. *radiodurans* was performed. The results reveal that CID boundaries have lower G+C content, and 78.3% of the CID boundaries (18 out of 23) are lower than the average level of chromosomal G+C contents. Moreover, the G+C content of CID boundaries is significantly lower than the interior (**Fig. S3**), which has also been reported in other species (46,47), suggesting a potential role of AT-rich DNA sequences in CID boundary formation. In view of the biased base composition of the CID boundary, we use the MEME software to search for conserved sequences (motifs) at these boundaries. Then the TOMTOM software was used to annotate and compare the motif sequences obtained, and it was found that some known transcription factor binding sites are significantly present in the boundary regions. For these motifs, visualization is present in **Figure 2C**, and detailed information is provided in **Table S2**. Some of them are also the known binding motifs for NAPs involved in transcriptional regulation, implying the potential influence of NAPs on the establishment or maintenance of local chromosome structure.

Furthermore, we explored the function of genes located at CID boundaries. Although some genes are unannotated, KEGG pathway annotation of the remaining 468 genes reveal that 80.3% of them are involved in metabolic processes, followed by genetic information processing (9.4%) and environmental information processing (7.5%), with only 2.8% of the genes associated with cellular processes (**Fig. 2D**). Further GO enrichment analysis of CID boundary genes show that the boundary genes are mainly enriched in the biological processes of metabolism and proton transmembrane transport. The gene products are predominantly located in the cytoplasm and ATPase complex, with molecular functions related to ATP binding, transport, and catalysis (**Fig. 2E**). These results reflect the important roles of CID boundary genes in the growth and metabolic processes of *D*. *radiodurans*.

**Figure 2.**
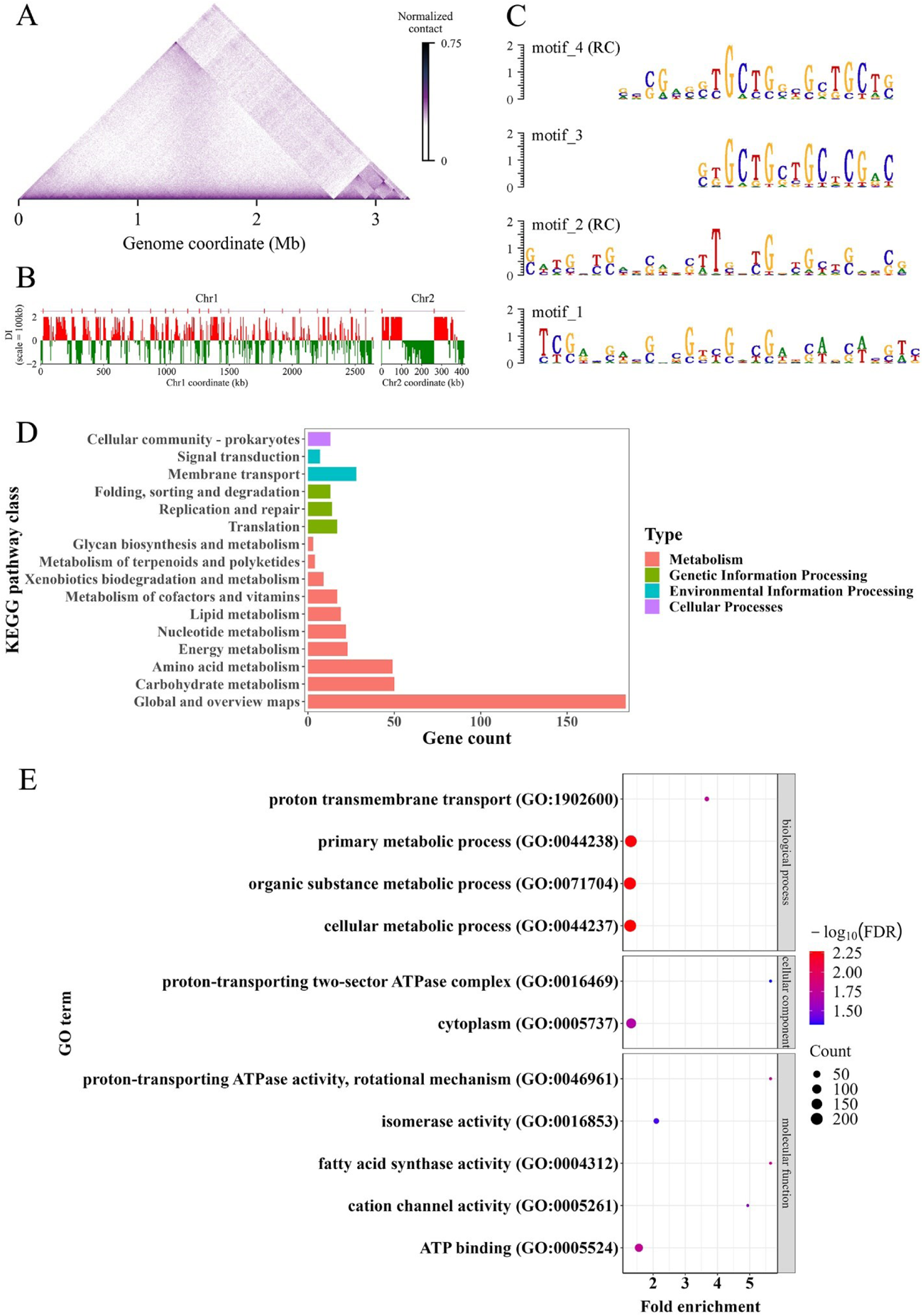
CID analysis of *D. radiodurans* chromosomes at 5 kb resolution. (A) Normalized contact map for the genome rotated 45° clockwise. (B) Domain boundaries characterized for wild-type condition using DI analysis performed at a scale of 100 kb. Downstream (red) and upstream (green) biases are indicated. Significant DI boundaries defining CIDs are annotated with red vertical lines above the panel. (C) Motifs at CID boundaries discovered through MEME software. RC represents the motifs discovered on the reverse complementary chain. (D) Kyoto Encyclopedia of Genes and Genomes (KEGG) pathway analysis of genes localized in CID boundaries. (E) Gene Ontology (GO) enrichment results of the CID boundary genes.

### Interplay between transcription and local chromosome structure

Previous studies have demonstrated that long and highly expressed transcripts in *B. subtilis* can drive local chromosome decompaction and promote spatial separation of gene flanking sequences, thereby reducing contact between DNA in neighboring domains to form CID boundaries (12). This phenomenon is also found in *D. radiodurans*. After obtaining gene expression profiles through transcriptome sequencing, we examined the lengths and transcriptional levels of genes at CID boundary and interior. **Figure 3A** shows that long genes with active transcription significantly exist at CID boundaries, which holds true for both UV irradiation condition and ΔDr_ebfC mutation.

Moreover, we investigated the relationship between transcription and short-range interactions. At a 5 kb resolution, we plotted the changes in interaction frequency and transcriptional level along the genome (**Fig. 3B**), revealing a significant positive correlation between the two sets of signals (Pearson correlation [PC] = 0.51). Notably, similar correlations have been observed in contact maps of other bacteria species generated by different laboratories using different enzymes and cross-linking conditions (6), indicating that such a correlation is a conservative feature. Additionally, the relationship between interaction frequency and genome distance was plotted according to gene expression level (**Fig. 3C**), which shows that the bins containing highly expressed genes have higher interaction frequencies with nearby bins, and the interaction frequency decays fast with the increase of linear distance. This suggests that the transcription process may limit the mobility of gene loci, resulting in reduced long-range interactions and increased short-range interactions.

**Figure 3.**
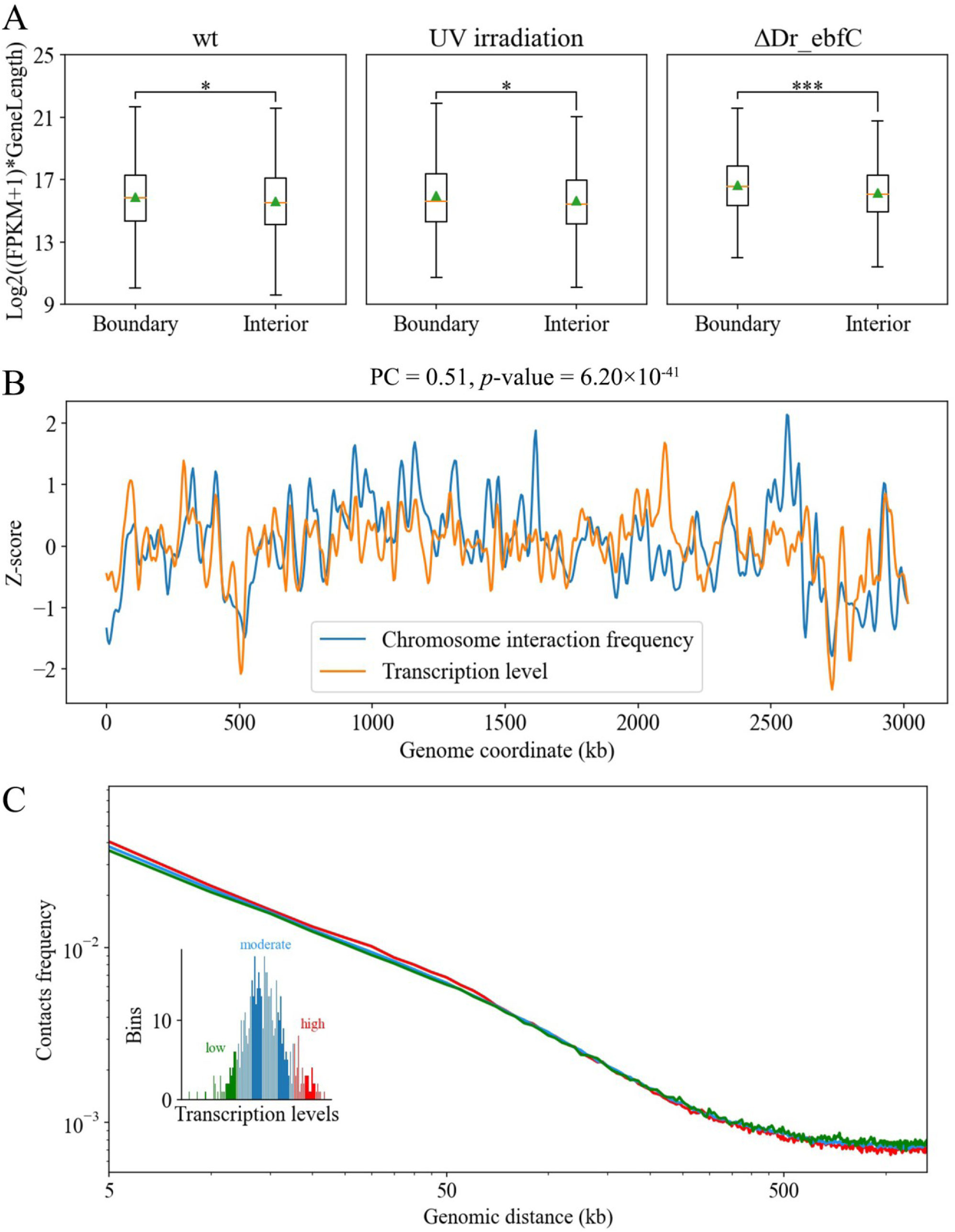
Correlation between transcription and local chromosome structure. (A) Box plots of CID boundary and interior gene expression distributions corresponding to different samples. The orange line indicates the median of the box. The green triangle indicates the average value of the box. ***, *p*-value < 0.001; *, *p*-value < 0.05. (B) Correlation between transcription level and short-range chromosome interaction frequency at 5 kb resolution. The abscissa denotes the location on the genome, while the ordinate represents the strength of the normalized signal value. The Pearson correlation (PC) between these two variables and the corresponding *p*-value are shown at the top of the graph. (C) Contact frequency as a function of genomic distance for genes categorized based on their expression levels: poorly expressed (green), moderately expressed (blue), or highly expressed (red). Inset: distribution of bin numbers for different transcription levels (at 5 kb resolution).

### UV irradiation alters chromosomal interaction regions

To investigate the impact of UV irradiation on the chromosomal structure of *D. radiodurans*, we performed 3C experiments on cells exposed to 750 J/m^2^ UV irradiation for 5 minutes. Viability assay reveals a survival rate of approximately 76.6% under this UV dosage (31), indicating sufficient live cells for 3C experiments. Irradiation stress often triggers responses in DNA recombination repair and base excision repair pathways. Analysis of normalized interaction matrix shows that UV irradiation led to weaker interaction (lighter color) along the main diagonal and stronger interaction (darker color) at other positions compared to the control group cells (**Fig. 4A, B**). To further visualize the interaction difference, ratio plot is used to show the interaction between each bin and its adjacent bins. The results indicate a reduction in short-range interactions and an increase in long-range interactions after UV irradiation (**Fig. 4C**). To confirm this phenomenon, we employed the Fit-Hi-C software to count significant interactions between pairs of bins at a 5 kb resolution across the whole genome. The results show that the interaction frequency of chromosome decrease with the increase of linear distance, with fewer significant short-range interactions (<450 kb) and more significant long-range interactions (>450 kb) after irradiation compared with control group cells (**Fig. 4D**). Furthermore, we identified 18 CIDs from the contact map of irradiated cells, ranging in size from 10 kb to 335 kb, with an average size of 139 kb. Five CIDs are conserved, and the larger CIDs observed in irradiated cells seem being formed by the fusion of several smaller CIDs present in the control group cells (**Fig. S4**).

Meanwhile, we performed transcriptome analysis on the irradiated cells and identified 750 differentially expressed genes (DEGs) with a fold change threshold of 2. Among these, upregulated genes (544) account for 72.5% and are significantly enriched in DNA recombination repair, SOS response, and translation processes (31). To explore the relationship between DEGs, particularly upregulated genes, and chromosomal structural changes, we analyzed the distribution of differentially upregulated genes on *D. radiodurans* Chr1 using a sliding step of 5 kb and a window size of 100 kb. The results, as shown in **Figure 4E, 4F** and **4G**, reveal that the distribution of differentially upregulated genes along the genome is not random but significantly enriched near certain CID boundaries. These special CID boundaries are mainly divided into two categories: the first consists of CID boundaries that are present in the control group cells but disappear after irradiation (e.g., ID_4, 5, 16, 17, 18, 19, 20, 21); the second comprises the conserved CID boundaries (e.g., ID_1, 2, 3, 9, 11, 12) in both irradiated and control group cells. Furthermore, GO enrichment results of CID boundary genes specific to the control group and the irradiation group were compared and analyzed (**Fig. S5**), and it was found that the function of boundary genes changed significantly. After UV irradiation, the specific CID boundary genes are mainly involved in the translation and ribosome assembly processes, and the protein products constitute the respiratory chain complex, ribosome subunit structure and other components, and perform the functions of ribosome structure maintenance and rRNA binding. The damage repair pathways of *D. radiodurans* in response to UV irradiation require the substantial synthesis of relevant proteins and the generation of ATPs through respiration. These results collectively suggest that *D. radiodurans* employs a coordinated response of chromosomal organization and transcriptional regulation to cope with UV irradiation stress and minimize damage to the bacterial cell.

**Figure 4.**
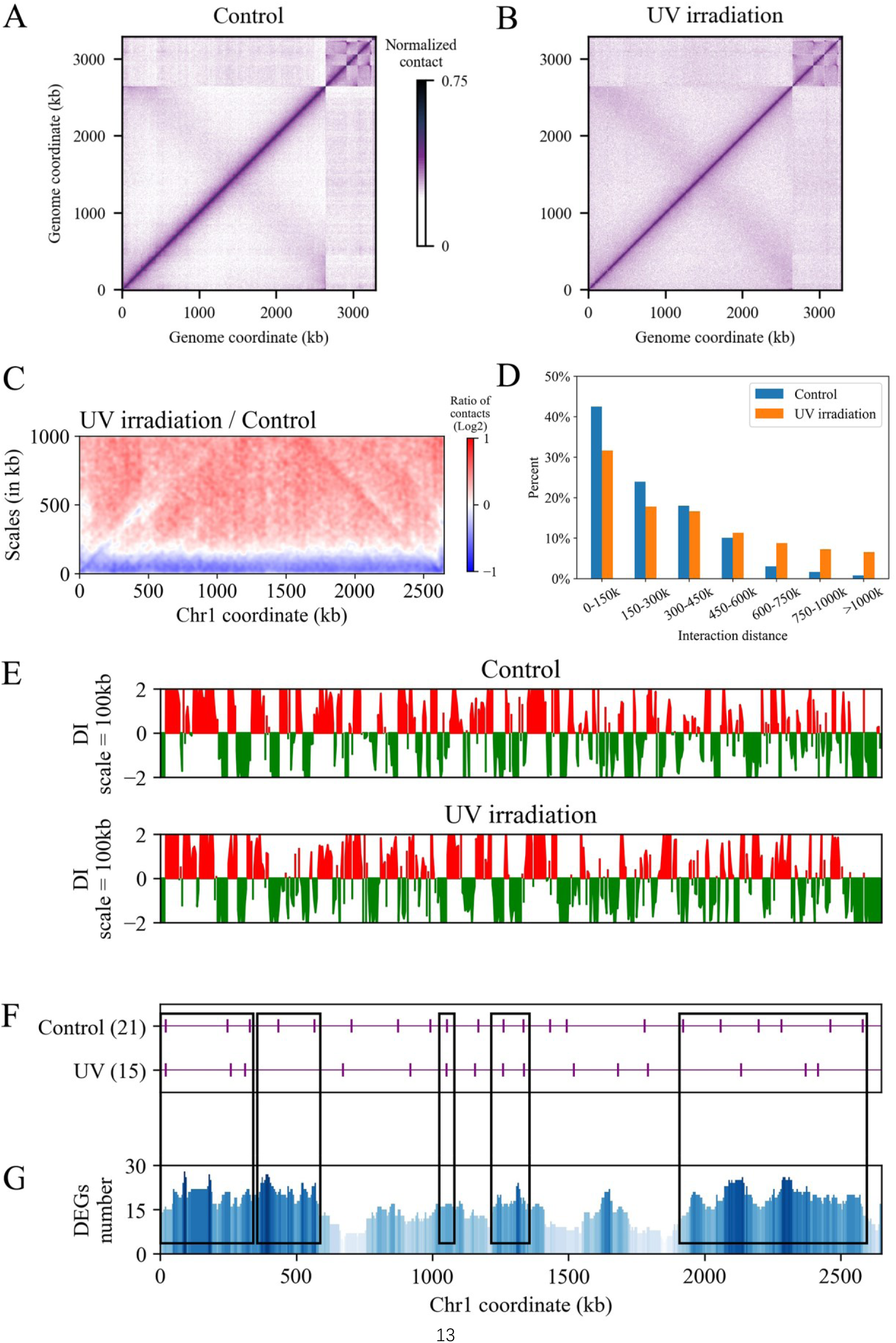
Effect of UV irradiation on chromosome contacts and CID boundaries. (A), (B) Normalized contact maps for the control group and UV irradiation condition, respectively. (C) Ratio (UV / Control) plot of the contact signals for each bin along Chr1. The *x-*axis indicates the position of the bin along the genome. The *y-*axis indicates the distance from the bin. A decrease or increase in contacts at the UV condition compared with the control is represented with a blue or red color, respectively. White color indicates no difference between the two conditions. (D) Proportion of significant *cis*-interactions of Chr1. The significant *cis*-interactions are classified according to different distance ranges. The interaction distance is calculated based on the genomic coordinate in the circular genome. If the distance between the significant interacting loci exceeds half of the length of Chr1, the shorter arc length is considered as the interaction distance. The *x*-axis is the interval of different interaction distances; the *y*-axis is the proportion of significant interactions within the distance interval to the total significant interactions. (E) DI analysis (100 kb) of Chr1 at Control and UV conditions, respectively. (F) CID boundaries along the Chr1 are marked by purple vertical lines. (G) Distribution of upregulated DEGs in Chr1. The black boxes across the subplots F and G represent the CID boundaries with enriched upregulated DEGs.

### The role of Dr_ebfC protein in chromosome organization

The Dr_ebfC protein in *D. radiodurans*, encoded by the *dr_0199* gene, was initially characterized as a NAP with DNA-protective and histone-like properties, safeguarding the DNA from damage (28). Its homologs are almost universal in all bacteria, showing a high degree of sequence conservation (**Fig. S6**). Sequence analysis reveal that the Dr_ebfC protein contains domains responsible for PPI and participates in DNA binding through the formation of a “tweezer-like” structure (29). To study the role of this protein in chromosomal organization, we constructed the *dr_0199* deletion mutant in *D. radiodurans*. The mutant strain is viable, but compared with wild-type cells, the growth rate is moderately inhibited and the cell density during the stationary phase is lower than that of the wild-type (**Fig. 5A**).

Through RNA-seq analysis of the mutant strain, we identified a total of 1077 DEGs, with 636 upregulated genes and 441 downregulated genes compared to the wild-type (**Fig. 5B**). GO and KEGG pathway analyses indicate that the upregulated genes in the mutant are mainly associated with environmental stress response, DNA recombination, regulation of metabolic activities, catalytic activities, and DNA binding processes (**Fig. S7A**, **B**). Notably, several DNA damage repair-related genes (e.g., *ddrA*, *ddrB*, *ddrC*, *ddrD*, *pprA*) regulated by PprI/DdrO proteins show significantly increased expression levels in the mutant (**Table S3**). This observation suggests that the absence of the Dr_ebfC protein might lead to cellular stress, implying a potential role of this NAP in DNA protection. The PprI/DdrO-mediated regulatory system has been recognized as a unique transcriptional inhibition and removal system of *D. radiodurans* in response to DNA damage. Under environmental stress conditions, PprI, a metal-dependent protease, is activated to specifically cleave the C-terminal region of the DdrO, abolishing its DNA binding activity, and resulting in the expression of DNA damage repair genes that are inhibited by the DdrO (48). Enrichment analysis of the downregulated genes in the mutant reveal their involvement in vital cellular processes closely related to respiration, translation, RNA metabolism, and electron transfer (**Fig. S7C**, D), corroborating the observed slow growth phenotype of the mutant. Additionally, genes encoding oxidative stress resistance proteins, such as catalase (*dr_1998*, *dr_a0259*, *dr_a0146*) and superoxide dismutase (*dr_1279*, *dr_1546*), show significant down-regulation in the mutant (**Table S4**). Among the three known catalases in *D. radiodurans*, DR1998 is the most important one, exhibiting a high activity of 68,800 U/mg and serving as a more effective scavenger of hydrogen peroxide compared to the other two enzymes (49). The transcription levels of *pdxT* and *pdxS* genes, encoding pyridoxine biosynthesis proteins, are also significantly decreased by more than threefold in the mutant strain. The enzymes of pyridoxal 5′-phosphate biosynthesis participate in the biosynthesis of vitamin B6, acting as efficient quencher of singlet oxygen and potential antioxidant (50). The down-regulation of these genes indicates a reduced resistance of ΔDr_ebfC mutant to oxidative damage, which is further confirmed by the hydrogen peroxide sensitivity assay of the mutant strain, as shown in **Figure 5C**.

The 3C-seq analysis of wild-type and ΔDr_ebfC cells show that the interaction matrices seem similar overall (**Fig. 6A**). However, the plot of interaction frequency as a function of distance exhibits a significant reduction in short-range interactions in the mutant compared to the wild-type, extending up to approximately 100 kb (**Fig. 6B**). Moreover, we used DI to calculate the degree of interaction preference of each bin on Chr1. Although the overall correlation of interaction preference between wild-type and mutant is relatively high (*r* = 0.63, *p* = 1.6 × 10^-60^), indicating some CID boundaries remain stable, a complete loss of 7 CID boundaries around the region of 750-1750 kb (near the *ter* region) in the ΔDr_ebfC mutant is observed (**Fig. 6C**). Further analysis using Chromosight software to detect chromosomal loop structures reveals a reduction in the number and strength of loop structures in this region of Chr1 in the mutant (**Fig. S8A**, **B**). More visually, the reconstructed structure of Chr1 shows that the ΔDr_ebfC mutant has more overall bending than the wild-type (**Fig. 6E**). There is also a significant difference in chromosome structure near the *ter* region where the CID boundaries are lost in the mutant (**Fig. 6F**). To quantitatively characterize the changes in spatial structure, we calculated the gyration radius for the reconstructed structure models of the wild-type and mutant chromosomes. The results show that the gyration radius is significantly smaller in the mutant than in the wild-type (**Fig. 6G**), suggesting a more compact 3D chromosome structure for the mutant strain. These analyses indicate that Dr_ebfC protein promotes local short-range interactions in the genome, and may play an important role in maintaining interactions between loci in the *ter* region.

To explore whether the 3D structure changes affect gene expression, we analyzed the DEGs corresponding to this region. We found significant difference in transcription levels between the mutant and wild-type on Chr1, where about 30% of the genes show significant changes (**Fig. 6D**). Specifically, in the region of 1480-1523 kb, CID boundary exists only in the wild-type and disappears in the mutant (**Fig. 6C**). The expression of a group of genes corresponding to this region is significantly reduced (**Table 1**), and the products encoded are mainly subunits constituting NADH-quinone oxidoreductase which involves in the electron transport chain and plays a crucial role in cellular respiration and energy production. Considering the slower growth rate of the mutant, we speculated that the DR_ebfC protein may influence the expression of related genes by altering the local chromosome conformation, thereby regulating the physiological state of the bacterium.

**Figure 5.**
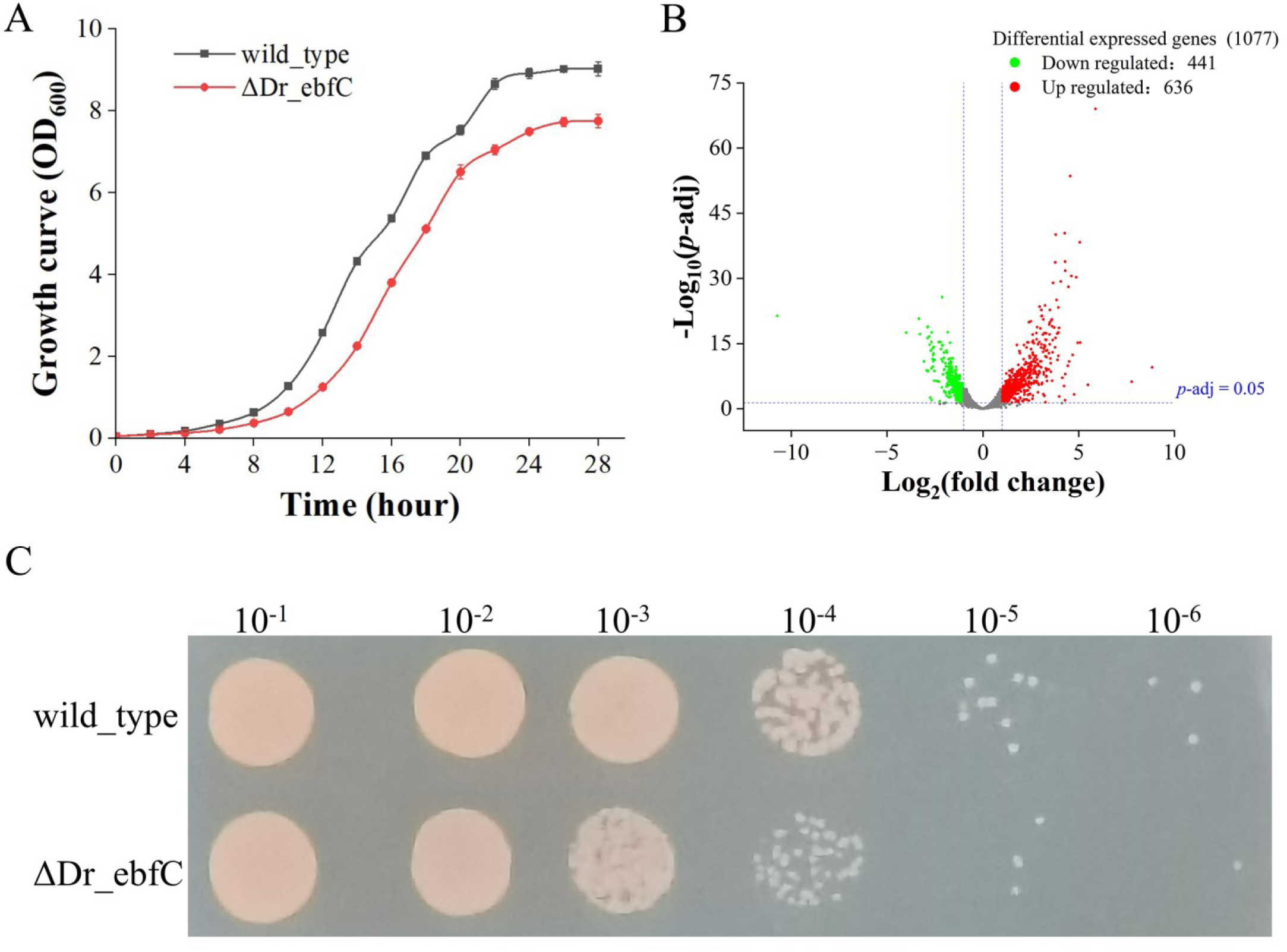
Comparative analysis of phenotypes and transcriptional profiles between wild-type and ΔDr_ebfC mutant strains. (A) Growth rate of wild-type and ΔDr_ebfC strain in TGY. (B) Volcano plot of DEGs in the ΔDr_ebfC strain compared with *D. radiodurans* wild type. Genes with an adjusted *p* < 0.05 are assigned as DEGs. (C) Susceptibility experiment of wild-type and ΔDr_ebfC strains of *D. radiodurans* to H2O2. After incubating in a 50 mM hydrogen peroxide solution for 30 minutes, the diluted bacterial suspension was spotted onto TGY agar plates. The values above the picture represent the dilution ratio of the bacterial suspension.

**Figure 6.**
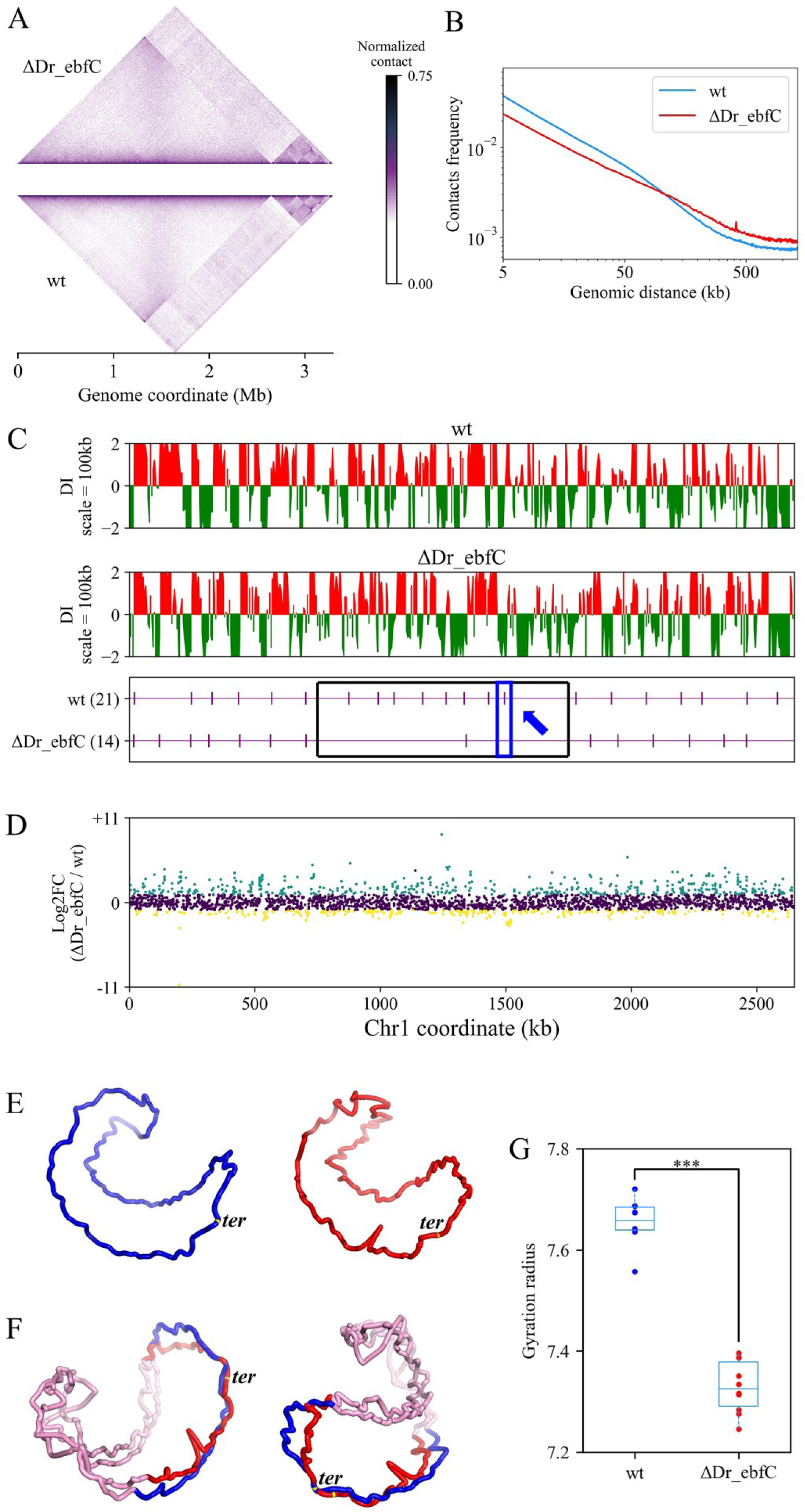
The role of Dr_ebfC protein in *D. radiodurans* chromosome organization. (A) Symmetric halves of the normalized contact maps of ΔDr_ebfC mutant (top) and wild-type (bottom) cells are aligned after clockwise rotation. (B) Plot of chromosome interaction frequency as a function of genomic distance. (C) CID distribution on Chr1 at 5 kb resolution of wild-type and ΔDr_ebfC strains, respectively. The purple vertical lines below represent CID boundaries along the Chr1. The CID boundaries lost in the ΔDr_ebfC strain are highlighted by the black box. The area with boundary loss and gene expression changes is marked by the blue arrow. (D) The distribution of DEGs of wild-type and ΔDr_ebfC strains on Chr1. Cyan and yellow dots represent upregulated and downregulated DEGs (fold change > 2 and FDR < 0.05), respectively. Purple dots represent genes with no significant change in expression. (E) The models of Chr1 for wild-type (blue) and ΔDr_ebfC mutant (red). (F) Comparison of the chromosome 3D structure near the *ter* region with missing CID boundaries (blue for wild-type and red for ΔDr_ebfC mutant, shown in two different perspectives). Other regions of the chromosome are colored in pink. (G) Boxplot for the gyration radius distribution of the 3D chromosome modes of wild-type and ΔDr_ebfC strains. The difference is significant (***, *p*-value < 0.001).

**Table 1.** A part of genes in the purple arrow region of Chr1 in Fig. 6C.

| Gene ID | Position (bp) |  | Product | Differential expression |
| --- | --- | --- | --- | --- |
| DR_RS07640 | 1506995 | 1508473 | NADH-quinone oxidoreductase subunit N | down (log<sub>2</sub>Foldchange = -2.72) |
| DR_RS07645 | 1508474 | 1509898 | NADH-quinone oxidoreductase subunit M | down (log<sub>2</sub>Foldchange = -2.89) |
| DR_RS07650 | 1509948 | 1511879 | NADH-quinone oxidoreductase subunit L | down (log<sub>2</sub>Foldchange = -2.80) |
| DR_RS07655 | 1511886 | 1512188 | NADH-quinone oxidoreductase subunit NuoK | down (log<sub>2</sub>Foldchange = -2.67) |
| DR_RS07660 | 1512190 | 1512807 | NADH-quinone oxidoreductase subunit J | down (log<sub>2</sub>Foldchange = -2.77) |
| DR_RS07665 | 1512804 | 1513340 | NADH-quinone oxidoreductase subunit NuoI | down (log<sub>2</sub>Foldchange = -2.60) |
| DR_RS07670 | 1513512 | 1514705 | NADH-quinone oxidoreductase subunit NuoH | down (log<sub>2</sub>Foldchange = -2.61) |
| DR_RS07675 | 1514705 | 1516897 | NADH-quinone oxidoreductase subunit NuoG | down (log<sub>2</sub>Foldchange = -2.52) |
| DR_RS07680 | 1517019 | 1518353 | NADH-quinone oxidoreductase subunit NuoF | down (log<sub>2</sub>Foldchange = -2.86) |
| DR_RS07685 | 1518350 | 1518973 | NADH-quinone oxidoreductase subunit NuoE | down (log<sub>2</sub>Foldchange = -3.08) |
| DR_RS07690 | 1518940 | 1519416 | hypothetical protein | down (log<sub>2</sub>Foldchange = -2.94) |
| DR_RS07695 | 1519446 | 1520651 | NADH dehydrogenase (quinone) subunit D | down (log<sub>2</sub>Foldchange = -2.89) |
| DR_RS07700 | 1520648 | 1521322 | NADH-quinone oxidoreductase subunit C | down (log<sub>2</sub>Foldchange = -2.54) |
| DR_RS07705 | 1521319 | 1521864 | NADH-quinone oxidoreductase subunit B | down (log<sub>2</sub>Foldchange = -2.69) |
| DR_RS07710 | 1521971 | 1522306 | NADH-quinone oxidoreductase subunit A | down (log<sub>2</sub>Foldchange = -2.30) |

## Discussion

Owing to the maturation of methodology, 3C/Hi-C techniques have been applied in more and more bacteria to investigate their genome organization and gene regulation over the past decade. In this study, we explored the 3D chromosomal architecture of the extremophile *D. radiodurans*, and investigated the effects of UV irradiation on genome organization. Integrating transcriptome data, we also elucidated the role of Dr_ebfC as a NAP in gene regulation and chromosomal organization.

We present genome-wide DNA interactions for all the four replicons of *D. radiodurans* at a 5 kb resolution. Among them, contact maps of the two chromosomes (Chr1, Chr2) show similar interaction domains, but different inter-arm interactions patterns (**Fig. S1**). Chr1 displays secondary diagonal interactions, while Chr2 does not, indicating different folding states for these two chromosomes. Visibly, it seems that the interactions between Chr2 and the two plasmids are stronger than those between other replicons (**Fig. 1A**). This is demonstrated by quantitative analysis of interaction frequencies between replicons, and the differences are statistically significant (**Fig. S9**). This phenomenon is different from the highly segmented *B. burgdorferi* genome. In *B. burgdorferi*, interactions between ∼18 plasmids are significantly higher than those between plasmids and chromosome, and the interactions between circular plasmids and linear plasmids exhibit marked differences (51). It is worth noting that although the genomes of both *D. radiodurans* and *B. burgdorferi* are multi-copied, the number of copies of each replicon in *B. burgdorferi* does not exceed two, while the number of genome copies of *D. radiodurans* has never been reported to be less than four (52). We infer that the number and copy count of replicons may be factors influencing interactions between plasmids.

The mechanism of coordinated replication between chromosomes is of interest in other studied bacteria with multipartite genome. In *V. cholerae*, the Ori region of Chr2 strongly interacts with the *crtS* region on Chr1, while the terminus regions of both Chr1 and Chr2 also exhibit preferential contacts (3). This is consistent with the mechanism by which two dissimilarly sized circular chromosomes of *V. cholerae* coordinate replication initiation but terminate replication at the same time. In *Agrobacterium tumefaciens*, the Ori regions of all four replicons are bundled together, resulting in an interaction pattern between the replicons that resembles a butterfly shape (9). To show the interactions between the two chromosomes of *D. radiodurans*, we positioned the Ori region at the center of each chromosome (**Fig. S10**). Unfortunately, we did not observe preferential contact between the origins or other regions on Chr1 and Chr2. It has been proposed that having multiple replicons enables faster genome replication and gives these bacteria the advantage of adapting quickly when switching hosts or environments (53). *D. radiodurans* also has multiple replicons and exhibits strong environmental adaptability, warranting further investigation into how these replicons coordinate and maintain genome stability within the bacterium.

The structural maintenance of chromosome (SMC) protein complex plays a pivotal role in shaping the global architecture of chromosomes, and extensive literature has elucidated the functions of numerous bacterial SMC proteins and their homologs (4,6,54). In this study, although the *smc* gene was not knocked out, transcriptome analysis of UV irradiation condition and ΔDr_ebfC strain found that there were no significant differences in *smc* gene expression compared to the wild type. Notably, the absence of a homolog of the ScpA protein (55), one of the subunits that constitute the SMC complex in *D. radiodurans*, possibly contributes to the faint secondary diagonal of the contact map. Previous studies have shown that the SMC-ScpAB holo-complex can tether the two arms of the chromosome together and promote chromosome compaction and separation (54).

*D. radiodurans* has long been an ideal model organism for studying DNA damage repair processes following irradiation. We observed that the frequency of short-range interaction decreased after UV irradiation. DI analysis of the chromosome interaction data revealed the fusion of CIDs through eliminating boundaries. CIDs reduced and transitioned from smaller to larger domains in the irradiated cells. Combined with transcriptome analysis, we found that the number of upregulated genes was more than twice that of downregulated genes, with the upregulated genes predominantly located near specific CID boundaries in the genome. CID boundaries often corresponded to long and active transcripts in *D. radiodurans*, which has also been demonstrated in *B. subtilis* (12). We conjecture that highly expressed long genes may more readily influence the preference of local interaction, affecting contacts between neighboring structures.

We investigated the role of Dr_ebfC as a NAP in *D. radiodurans* using knockout inactivation strategy. Phenotypic experiments reveal that the ΔDr_ebfC mutant is viable but exhibits slower growth and increased sensitivity to oxidative stress compared to the wild-type, which align well with previous findings conducted by Wang et al (28). Transcriptomic analysis of the ΔDr_ebfC mutant indicates down-regulation of genes involved in respiration and translation, providing molecular insights into the impact of Dr_ebfC as a regulatory factor on bacterial physiological activities. The Dr_ebfC protein has similar properties to other NAPs and can regulate gene expression by controlling chromosome topology (56–58). We identified a region where changes in chromosomal structure resulted in the down-regulation of a group of genes, leading to a slowdown in energy production and the slow growth phenotype of the mutant. While further studies are needed to elucidate the causal relationship between structural change and gene regulation, the exploratory results in this work have expanded our understanding on the chromosome behaviors of this extremophilic microorganism in response to environmental stresses.

## Funding

This work was supported by the National Natural Science Foundation of China (Grant 31971184).

## Notes

### Competing Interest Statement

The authors have declared no competing interest.

## References

1. Dame, R.T., Rashid, F.-Z.M. and Grainger, D.C. (2020) Chromosome organization in bacteria: mechanistic insights into genome structure and function. Nat. Rev. Genet., 21, 227–242.

2. Le, T.B., Imakaev, M.V., Mirny, L.A. and Laub, M.T. (2013) High-resolution mapping of the spatial organization of a bacterial chromosome. Science, 342, 731–734.

3. Val, M.E., Marbouty, M., de Lemos Martins, F., Kennedy, S.P., Kemble, H., Bland, M.J., Possoz, C., Koszul, R., Skovgaard, O. and Mazel, D. (2016) A checkpoint control orchestrates the replication of the two chromosomes of *Vibrio cholerae*. Sci. Adv., 2, e1501914.

4. Tran, N.T., Laub, M.T. and Le, T.B.K. (2017) SMC progressively aligns chromosomal arms in *Caulobacter crescentus* but is antagonized by convergent transcription. Cell Rep., 20, 2057–2071.

5. Trussart, M., Yus, E., Martinez, S., Bau, D., Tahara, Y.O., Pengo, T., Widjaja, M., Kretschmer, S., Swoger, J., Djordjevic, S. et al. (2017) Defined chromosome structure in the genome-reduced bacterium *Mycoplasma pneumoniae*. Nat. Commun., 8, 14665.

6. Lioy, V.S., Cournac, A., Marbouty, M., Duigou, S., Mozziconacci, J., Espeli, O., Boccard, F. and Koszul, R. (2018) Multiscale structuring of the *E. coli* chromosome by nucleoid-associated and condensin proteins. Cell, 172, 771–783.

7. Lioy, V.S., Junier, I., Lagage, V., Vallet, I. and Boccard, F. (2020) Distinct Activities of Bacterial Condensins for Chromosome Management in *Pseudomonas aeruginosa*. Cell Rep., 33, 108344.

8. Conin, B., Billault-Chaumartin, I., El Sayyed, H., Quenech’Du, N., Cockram, C., Koszul, R. and Espeli, O. (2022) Extended sister-chromosome catenation leads to massive reorganization of the *E. coli* genome. Nucleic Acids Res., 50, 2635–2650.

9. Ren, Z., Liao, Q., Karaboja, X., Barton, I.S., Schantz, E.G., Mejia-Santana, A., Fuqua, C. and Wang, X. (2022) Conformation and dynamic interactions of the multipartite genome in *Agrobacterium tumefaciens*. Proc. Natl. Acad. Sci. U.S.A., 119, e2115854119.

10. Dixon, J.R., Selvaraj, S., Yue, F., Kim, A., Li, Y., Shen, Y., Hu, M., Liu, J.S. and Ren, B. (2012) Topological domains in mammalian genomes identified by analysis of chromatin interactions. Nature, 485, 376–380.

11. Nora, E.P., Lajoie, B.R., Schulz, E.G., Giorgetti, L., Okamoto, I., Servant, N., Piolot, T., van Berkum, N.L., Meisig, J., Sedat, J., et al. (2012) Spatial partitioning of the regulatory landscape of the X-inactivation centre. Nature, 485, 381–385.

12. Le, T.B. and Laub, M.T. (2016) Transcription rate and transcript length drive formation of chromosomal interaction domain boundaries. EMBO J., 35, 1582–1595.

13. Amemiya, H.M., Schroeder, J. and Freddolino, P.L. (2021) Nucleoid-associated proteins shape chromatin structure and transcriptional regulation across the bacterial kingdom. Transcription, 12, 182–218.

14. Le Berre, D., Reverchon, S., Muskhelishvili, G. and Nasser, W. (2022) Relationship between the chromosome structural dynamics and gene expression – a chicken and egg dilemma? Microorganisms, 10, 846.

15. Dugar, G., Hofmann, A., Heermann, D.W. and Hamoen, L.W. (2022) A chromosomal loop anchor mediates bacterial genome organization. Nat. Genet., 54, 194–201.

16. Slade, D. and Radman, M. (2011) Oxidative stress resistance in *Deinococcus radiodurans*. Microbiol. Mol. Biol. Rev., 75, 133–191.

17. Battista, J.R. (1997) Against all odds: the survival strategies of *Deinococcus radiodurans*. Annu. Rev. Microbiol., 51, 203–224.

18. Im, S., Joe, M., Kim, D., Park, D.H. and Lim, S. (2013) Transcriptome analysis of salt-stressed *Deinococcus radiodurans* and characterization of salt-sensitive mutants. Res. Microbiol., 164, 923–932.

19. Joe, M.H., Jung, S.W., Im, S.H., Lim, S.Y., Song, H.P., Kwon, O. and Kim, D.H. (2011) Genome-wide response of *Deinococcus radiodurans* on cadmium toxicity. J. Microbiol. Biotechnol., 21, 438–447.

20. Liu, Y., Zhou, J., Omelchenko, M.V., Beliaev, A.S., Venkateswaran, A., Stair, J., Wu, L., Thompson, D.K., Xu, D., Rogozin, I.B. et al. (2003) Transcriptome dynamics of *Deinococcus radiodurans* recovering from ionizing radiation. Proc. Natl. Acad. Sci. U.S.A., 100, 4191–4196.

21. Wu, Y., Chen, W., Zhao, Y., Xu, H. and Hua, Y. (2009) Involvement of RecG in H_2_O_2_-induced damage repair in *Deinococcus radiodurans*. Can. J. Microbiol., 55, 841–848.

22. Xue, D., Liu, W., Chen, Y., Liu, Y., Han, J., Geng, X., Li, J., Jiang, S., Zhou, Z., Zhang, W. et al. (2019) RNA-seq-based comparative transcriptome analysis highlights new features of the heat-stress response in the extremophilic bacterium *Deinococcus radiodurans*. Int. J. Mol. Sci., 20, 5603.

23. Passot, F.M., Nguyen, H.H., Dard-Dascot, C., Thermes, C., Servant, P., Espeli, O. and Sommer, S. (2015) Nucleoid organization in the radioresistant bacterium *Deinococcus radiodurans*. Mol. Microbiol., 97, 759–774.

24. Toueille, M., Mirabella, B., Guerin, P., Bouthier de la Tour, C., Boisnard, S., Nguyen, H.H., Blanchard, L., Servant, P., de Groot, A., Sommer, S., et al. (2012) A comparative proteomic approach to better define *Deinococcus* nucleoid specificities. J Proteomics, 75, 2588–2600.

25. Grove, A. and Wilkinson, S.P. (2005) Differential DNA binding and protection by dimeric and dodecameric forms of the ferritin homolog Dps from *Deinococcus radiodurans*. J. Mol. Biol., 347, 495–508.

26. Nguyen, H.H., de la Tour, C.B., Toueille, M., Vannier, F., Sommer, S. and Servant, P. (2009) The essential histone-like protein HU plays a major role in *Deinococcus radiodurans* nucleoid compaction. Mol. Microbiol., 73, 240–252.

27. Santos, S.P., Yang, Y., Rosa, M.T.G., Rodrigues, M.A.A., De La Tour, C.B., Sommer, S., Teixeira, M., Carrondo, M.A., Cloetens, P., Abreu, I.A., et al. (2019) The interplay between Mn and Fe in *Deinococcus radiodurans* triggers cellular protection during paraquat-induced oxidative stress. Sci. Rep., 9, 17217.

28. Wang, H., Wang, F., Hua, X., Ma, T., Chen, J., Xu, X., Wang, L., Tian, B. and Hua, Y. (2012) Genetic and biochemical characteristics of the histone-like protein DR0199 in *Deinococcus radiodurans*. Microbiology, 158, 936–943.

29. Cordeiro, T., Gontijo, M.T.P., Jorge, G.P. and Brocchi, M. (2022) EbfC/YbaB: a widely distributed nucleoid-associated protein in prokaryotes. Microorganisms, 10, 1945.

30. Babb, K., Bykowski, T., Riley, S.P., Miller, M.C., Demoll, E. and Stevenson, B. (2006) *Borrelia burgdorferi* EbfC, a novel, chromosomally encoded protein, binds specific DNA sequences adjacent to *erp* loci on the spirochete’s resident cp32 prophages. J. Bacteriol., 188, 4331–4339.

31. Zhang, C.Y., Qiu, Q.T. and Ma, B.G. (2022) Transcriptomic analysis of *Deinococcus radiodurans* during the early recovery stage from ultraviolet irradiation. Prog. Biochem. Biophys., 50, 1701–1715.

32. Cai, J., Pan, C., Zhao, Y., Xu, H., Tian, B., Wang, L. and Hua, Y. (2022) DRJAMM is involved in the oxidative resistance in *Deinococcus radiodurans*. Front. Microbiol., 12, 756867.

33. Kim, D., Langmead, B. and Salzberg, S.L. (2015) HISAT: a fast spliced aligner with low memory requirements. Nat. Methods, 12, 357–360.

34. Pertea, M., Pertea, G.M., Antonescu, C.M., Chang, T.C., Mendell, J.T. and Salzberg, S.L. (2015) StringTie enables improved reconstruction of a transcriptome from RNA-seq reads. Nat. Biotechnol., 33, 290–295.

35. Love, M.I., Huber, W. and Anders, S. (2014) Moderated estimation of fold change and dispersion for RNA-seq data with DESeq2. Genome Biol., 15, 550.

36. Martin, M. (2011) Cutadapt removes adapter sequences from high-throughput sequencing reads. EMBnet j., 17, 10–12.

37. Langmead, B. and Salzberg, S.L. (2012) Fast gapped-read alignment with Bowtie 2. Nat. Methods, 9, 357–359.

38. Cournac, A., Marie-Nelly, H., Marbouty, M., Koszul, R. and Mozziconacci, J. (2012) Normalization of a chromosomal contact map. BMC Genom., 13, 436.

39. Kaul, A., Bhattacharyya, S. and Ay, F. (2020) Identifying statistically significant chromatin contacts from Hi-C data with FitHiC2. Nat. Protoc., 15, 991–1012.

40. Matthey-Doret, C., Baudry, L., Breuer, A., Montagne, R., Guiglielmoni, N., Scolari, V., Jean, E., Campeas, A., Chanut, P.H., Oriol, E. et al. (2020) Computer vision for pattern detection in chromosome contact maps. Nat. Commun., 11, 5795.

41. Wang, X., Gu, W.C., Li, J. and Ma, B.G. (2023) EVRC: reconstruction of chromosome 3D structure models using Error-Vector Resultant algorithm with Clustering coefficient. Bioinformatics, 39, btad638.

42. White, O., Eisen, J.A., Heidelberg, J.F., Hickey, E.K., Peterson, J.D., Dodson, R.J., Haft, D.H., Gwinn, M.L., Nelson, W.C., Richardson, D.L. et al. (1999) Genome sequence of the radioresistant bacterium *Deinococcus radiodurans* R1. Science, 286, 1571–1577.

43. Tian, L., Wang, X., Hua, K. and Ma, B. (2019) Bacteria 3D genomics. Chin. Sci. Bull., 64, 1780–1790.

44. Xie, T., Fu, L.Y., Yang, Q.Y., Xiong, H., Xu, H., Ma, B.G. and Zhang, H.Y. (2015) Spatial features for *Escherichia coli* genome organization. BMC Genom., 16, 37.

45. Szklarczyk, D., Gable, A.L., Nastou, K.C., Lyon, D., Kirsch, R., Pyysalo, S., Doncheva, N.T., Legeay, M., Fang, T., Bork, P. et al. (2021) The STRING database in 2021: customizable protein-protein networks, and functional characterization of user-uploaded gene/measurement sets. Nucleic Acids Res., 49, D605–D612.

46. Deng, L., Zhao, Z., Liu, L., Zhong, Z., Xie, W., Zhou, F., Xu, W., Zhang, Y., Deng, Z. and Sun, Y. (2023) Dissection of 3D chromosome organization in *Streptomyces coelicolor* A3(2) leads to biosynthetic gene cluster overexpression. Proc. Natl. Acad. Sci. U.S.A., 120, e2222045120.

47. Huang, Y.F., Liu, L., Wang, F., Yuan, X.W., Chen, H.C. and Liu, Z.F. (2023) High-resolution 3D genome map of *Brucella* chromosomes in exponential and stationary phases. Microbiol. Spectr., 11, e0429022.

48. Eugenie, N., Zivanovic, Y., Lelandais, G., Coste, G., Bouthier de la Tour, C., Bentchikou, E., Servant, P. and Confalonieri, F. (2021) Characterization of the radiation desiccation response regulon of the radioresistant bacterium *Deinococcus radiodurans* by integrative genomic analyses. Cells, 10, 2536.

49. Gao, L., Zhou, Z., Chen, X., Zhang, W., Lin, M. and Chen, M. (2020) Comparative proteomics analysis reveals new features of the oxidative stress response in the polyextremophilic bacterium *Deinococcus radiodurans*. Microorganisms, 8, 451.

50. Ott, E., Kawaguchi, Y., Kolbl, D., Rabbow, E., Rettberg, P., Mora, M., Moissl-Eichinger, C., Weckwerth, W., Yamagishi, A. and Milojevic, T. (2020) Molecular repertoire of *Deinococcus radiodurans* after 1 year of exposure outside the International Space Station within the Tanpopo mission. Microbiome, 8, 150.

51. Ren, Z., Takacs, C.N., Brandão, H.B., Jacobs-Wagner, C. and Wang, X. (2023) Organization and replicon interactions within the highly segmented genome of *Borrelia burgdorferi*. PLoS Genet., 19, e1010857.

52. Cox, M.M. and Battista, J.R. (2005) *Deinococcus radiodurans* — the consummate survivor. Nat. Rev. Microbiol., 3, 882–892.

53. Misra, H.S., Maurya, G.K., Kota, S. and Charaka, V.K. (2018) Maintenance of multipartite genome system and its functional significance in bacteria. J. Genet., 97, 1013–1038.

54. Wang, X., Brandão, H.B., Le, T.B.K., Laub, M.T. and Rudner, D.Z. (2017) *Bacillus subtilis* SMC complexes juxtapose chromosome arms as they travel from origin to terminus. Science, 355, 524–527.

55. Bouthier de la Tour, C., Toueille, M., Jolivet, E., Nguyen, H.-H., Servant, P., Vannier, F. and Sommer, S. (2009) The *Deinococcus radiodurans* SMC protein is dispensable for cell viability yet plays a role in DNA folding. Extremophiles, 13, 827–837.

56. Dorman, C.J. (2014) Function of nucleoid-associated proteins in chromosome structuring and transcriptional regulation. J. Mol. Microbiol. Biotechnol., 24, 316–331.

57. Browning, D.F., Grainger, D.C. and Busby, S.J.W. (2010) Effects of nucleoid-associated proteins on bacterial chromosome structure and gene expression. Curr. Opin. Microbiol., 13, 773–780.

58. Dillon, S.C. and Dorman, C.J. (2010) Bacterial nucleoid-associated proteins, nucleoid structure and gene expression. Nat. Rev. Microbiol., 8, 185–195.

