## Supplementary tables and figures for "Spatial chromosome organization and adaptation of the radiation-resistant extremophile *Deinococcus radiodurans*"

**Table S1.** List of strains, plasmid, and primers used in this experiment

| Name | Relevant characteristics, or sequence | Source |
|---|---|---|
| Strains | | |
| *D. radiodurans* R1 | Wild-type strain ATCC13939 | Laboratory stock |
| ΔDr_ebfC | R1 but *dr_0199::kan* | This study |
| Plasmid | | |
| pRADK | *D. radiodurans* shuttle vector | Laboratory stock |
| Primers | | |
| Δ*dr_0199*-p1 | CACTTCCAGTTCGGCCACGTCC | This study |
| Δ*dr_0199*-p2 | ACAGACGGATCCGACCGTCAGTCTAGCGGCT | This study |
| Δ*dr_0199*-p3 | CATAGAAGCTTGTGTGACCGCGCTGGC | This study |
| Δ*dr_0199*-p4 | CTTGCTCACGGTCTGGCGG | This study |
| *kan*-p5 | CATAGAAGCTTCGTATTGTCGCCCTACATAT | This study |
| *kan*-p6 | ACAGACGGATCCTAGAAAAACTCATCG | This study |
| Δ*dr_0199_test*-F | GCTTCGGGCTTGATTTTCAGG | This study |
| Δ*dr_0199_test*-R | CGGCTGGCAAGATTCAGGAC | This study |

**Table S2.** Predicted motifs of transcription factors and NAPs in CID boundaries

| Identifier | Width | E-value | Matched motif |
|---|---|---|---|
| motif_1 | 29 | 6.00E-15 | MX000107 (OxyR), MX000207 (McbR), MX000095 (MhpR), MX000147 (MetJ), MX000177 (TreR) |
| motif_2 | 29 | 1.00E-12 | MX000152 (GlnG), MX000103 (ExsA), MX000102 (Dnr), MX000095 (MhpR), MX000002 (Anr), MX000186 (AlgZ), MX000178 (TorR), MX000111 (RpoN), MX000070 (SigE), MX000163 (Lrp), MX000171 (RhaS), MX000132 (FliA), MX000190 (FNRL), MX000192 (NnrR) |
| motif_3 | 15 | 4.50E-09 | MX000192 (NnrR), MX000095 (MhpR), MX000152 (GlnG), MX000177 (TreR), MX000007 (ArgR), MX000180 (HydG), MX000119 (CpxR), MX000111 (RpoN), MX000002 (Anr), MX000101 (AlgR), MX000021 (Spo0A), MX000190 (FNRL), MX000046 (ComA) |
| motif_4 | 21 | 1.90E-07 | MX000095 (MhpR), MX000152 (GlnG), MX000192 (NnrR), MX000042 (RegR), MX000033 (AlgU), MX000163 (Lrp), MX000030 (DegU), MX000164 (Lrp), MX000208 (PrrA), MX000186 (AlgZ), MX000180 (HydG), MX000034 (AlgU), MX000111 (RpoN), MX000107 (OxyR), MX000126 (FlhD2C2), MX000069 (SigE) |
| motif_5 | 20 | 1.90E-07 | MX000180 (HydG), MX000042 (RegR), MX000101 (AlgR), MX000152 (GlnG), MX000192 (NnrR), MX000095 (MhpR), MX000033 (AlgU), MX000111 (RpoN), MX000034 (AlgU), MX000124 (FadR), MX000046 (ComA), MX000115 (AraC), MX000208 (PrrA), MX000186 (AlgZ), MX000178 (TorR), MX000002 (Anr), MX000126 (FlhD2C2) |
| motif_6 | 20 | 2.00E-07 | MX000095 (MhpR), MX000177 (TreR), MX000001 (MexR), MX000002 (Anr), MX000183 (TrpR), MX000008 (GlpR), MX000171 (RhaS), MX000033 (AlgU), MX000208 (PrrA), MX000101 (AlgR), MX000126 (FlhD2C2), MX000070 (SigE), MX000172 (RhaR) |
| motif_7 | 26 | 1.30E-06 | MX000171 (RhaS), MX000101 (AlgR), MX000065 (SigD), MX000095 (MhpR), MX000137 (LacI), MX000148 (ModE), MX000115 (AraC), MX000203 (DevR), MX000034 (AlgU), MX000001 (MexR), MX000177 (TreR), MX000104 (FleQ), MX000208 (PrrA), MX000139 (MalT) |
| motif_8 | 29 | 5.40E-06 | MX000203 (DevR), MX000186 (AlgZ), MX000107 (OxyR), MX000006 (SpoIIID), MX000103 (ExsA), MX000195 (YsiA), MX000130 (GalS), MX000127 (FhlA), MX000193 (KdgR), MX000191 (PpsR) |
| motif_9 | 21 | 1.70E-05 | MX000025 (DinR/LexA), MX000021 (Spo0A), MX000151 (NhaR), MX000095 (MhpR), MX000070 (SigE), MX000097 (Mlc), MX000107 (OxyR), MX000183 (TrpR), MX000207 (McbR), MX000172 (RhaR), MX000147 (MetJ), MX000176 (YdiH) |

**Table S3.** List of genes belonging to the PprI/DdrO regulon in *D. radiodurans* previously described by Eugénie et al [1].

| Gene ID | $Log_2FC$ | *P*-adj | Direction | Locus tag | Function description |
|---|---|---|---|---|---|
| | | | Replication, recombination, and repair (17) | | |
| DR_RS00525 | 0.76 | 3.29E-02 | ns | DR_0100 | single-stranded DNA-binding protein SSB |
| DR_RS02185 | 3.85 | 9.58E-26 | up | DR_0423 | single-stranded DNA-binding protein DdrA |
| DR_RS00370 | 3.05 | 1.95E-23 | up | DR_0070 | single-stranded DNA-binding protein DdrB |
| DR_RS00015 | 3.10 | 4.24E-22 | up | DR_0003 | DNA damage response protein DdrC |
| DR_RS01690 | 2.92 | 1.49E-10 | up | DR_0326 | DNA damage response protein DdrD |
| DR_RS09060 | -0.80 | 1.02E-02 | ns | DR_1771 | excinuclease ABC subunit UvrA |
| DR_RS11695 | -0.35 | 1.05E-01 | ns | DR_2275 | excinuclease ABC subunit UvrB |
| DR_RS12030 | 0.55 | 1.00E-01 | ns | DR_2340 | recombinase RecA |
| DR_RS09795 | -0.41 | 5.79E-02 | ns | DR_1913 | DNA gyrase subunit A |
| DR_RS04680 | -0.96 | 5.67E-04 | ns | DR_0906 | type IIA DNA topoisomerase subunit B |
| DR_RS03100 | 0.65 | 2.31E-02 | ns | DR_0596 | Holliday junction branch migration DNA helicase RuvB |
| DR_RS05355 | 2.09 | 6.85E-10 | up | DR_1039 | DNA mismatch repair protein MutS |
| DR_RS09810 | 0.25 | 4.26E-01 | ns | DR_1916 | ATP-dependent DNA helicase RecG |
| DR_RS09740 | 2.12 | 1.68E-09 | up | DR_1902 | ATP-dependent RecD-like DNA helicase |
| DR_RS10620 | -0.20 | 4.14E-01 | ns | DR_2069 | NAD-dependent DNA ligase LigA |
| DR_RS12020 | 1.80 | 5.56E-07 | up | DR_2338 | CinA family mononucleotide deamidase-related protein |
| DR_RS15335 | 1.30 | 5.05E-07 | up | DR_A0346 | DNA repair protein PprA |
| | | | Translation and post-translational modification (4) | | |
| DR_RS00725 | 0.17 | 6.45E-01 | ns | DR_0139 | GTPase HflX |
| DR_RS11140 | 0.24 | 3.20E-01 | ns | DR_2174 | leucine--tRNA ligase |
| DR_RS11600 | 2.45 | 1.52E-05 | up | DR_2255 | GNAT family N-acetyltransferase |
| DR_RS12555 | 0.04 | 8.97E-01 | ns | DR_2441 | GNAT family N-acetyltransferase |
| | | | Metabolism and metabolic transport (5) | | |
| DR_RS01110 | -0.24 | 4.21E-01 | ns | DR_0217 | sulfurtransferase |
| DR_RS02900 | -0.50 | 1.74E-01 | ns | DR_0561 | extracellular solute-binding protein |
| DR_RS06680 | 1.51 | 2.01E-06 | up | DR_1297 | DUF808 domain-containing protein |
| DR_RS11605 | 0.27 | 3.53E-01 | ns | DR_2256 | transketolase |
| DR_RS14970 | -1.68 | 5.23E-09 | down | DR_A0275 | cytochrome c |
| | | | Unknown function (7) | | |
| DR_RS01120 | -0.60 | 5.14E-02 | ns | DR_0219 | hypothetical protein |
| DR_RS03565 | 0.58 | 1.97E-02 | ns | DR_0685 | DUF11 domain-containing protein |
| DR_RS05905 | 2.17 | 2.65E-08 | up | DR_1143 | hypothetical protein |
| DR_RS08030 | -1.23 | 3.02E-04 | down | DR_1571 | ABC transporter substrate-binding protein |
| DR_RS11135 | 1.60 | 6.34E-06 | up | DR_2173 | DUF1963 domain-containing protein |
| DR_RS14425 | 0.46 | 1.31E-01 | ns | DR_A0165 | hypothetical protein |
| DR_RS16300 | 1.06 | 9.40E-03 | up | DR_C0023 | hypothetical protein |
| | | | Ambiguous (3) | | |
| DR_RS16270 | 3.62 | 1.09E-19 | up | DR_C0017 | Tn3 family transposase |
| DR_RS06675 | 0.68 | 0.004257 | ns | DR_1296 | IS5-like element ISDra5 family transposase |
| DR_RS16355 | 0.50 | 0.037748 | ns | DR_C0033 | IS5-like element ISDra5 family transposase |

Note: The three columns of $Log_2FC$, *P*-adj, and Direction are derived from the differential expression analysis of mutant and wild-type cells. Up and down indicate |fold change| > 2 and *P*-adj < 0.05, respectively. Ns indicates |fold change| < 2 or *P*-adj > 0.05.

**Table S4.** Genes involved in oxidative resistance

| Gene ID | $Log_2FC$ | *P*-adj | Gene name | Locus tag | Function description |
|---|---|---|---|---|---|
| DR_RS10235 | -2.08 | 1.27E-08 | *katA* | DR_1998 | catalase |
| DR_RS14880 | -1.28 | 1.46E-04 | *katE* | DR_A0259 | catalase |
| DR_RS14330 | -1.44 | 7.11E-06 | | DR_A0146 | catalase family protein |
| DR_RS06595 | -1.53 | 7.78E-07 | *sodA* | DR_1279 | superoxide dismutase [Mn] |
| DR_RS07910 | -1.46 | 2.39E-07 | | DR_1546 | superoxide dismutase family protein |
| DR_RS07020 | -1.80 | 3.94E-06 | *pdxT* | DR_1366 | pyridoxal 5'-phosphate synthase glutaminase subunit PdxT |
| DR_RS07025 | -1.86 | 1.63E-06 | *pdxS* | DR_1367 | pyridoxal 5'-phosphate synthase lyase subunit PdxS |

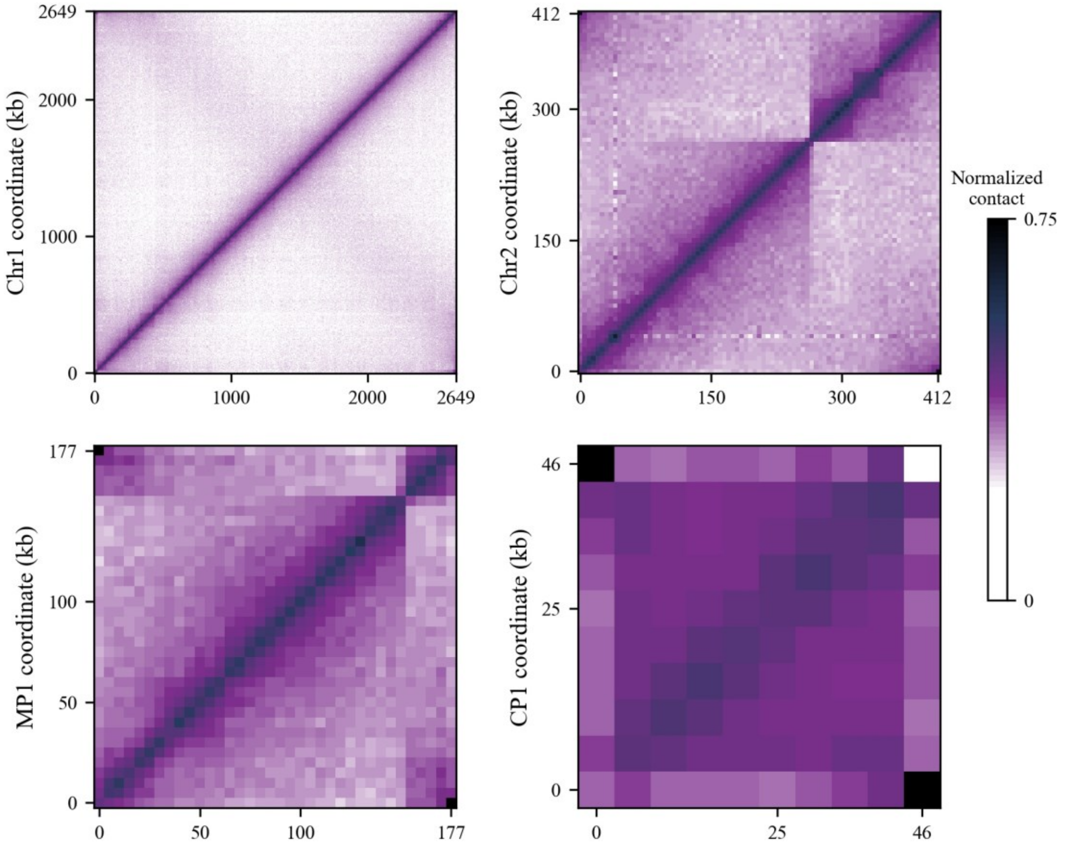


**Figure S1.** Chromosome contact map of the four replicons of *D. radiodurans.*

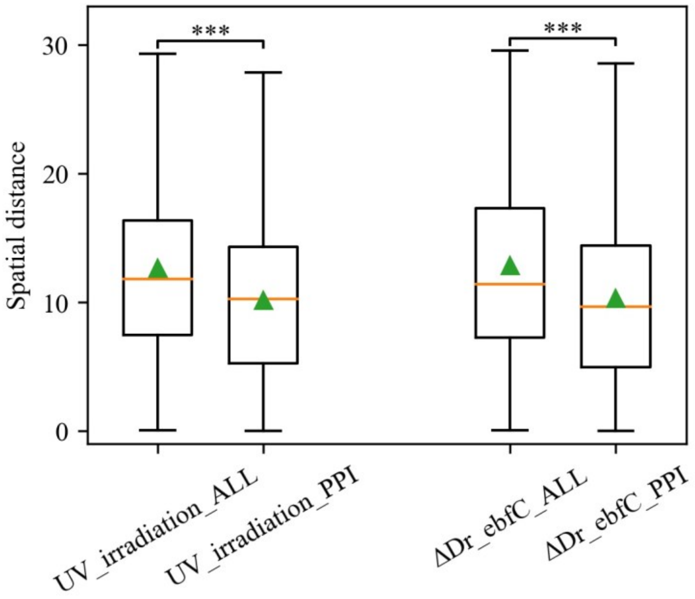


**Figure S2.** Boxplot of the spatial distance distribution between bin pairs in the 3D mode. The two groups (ALL vs. PPI) on the *x*-axis correspond to the distribution of spatial distance between all bins (ALL) and between the bins containing protein-protein interaction (PPI), respectively. The orange line indicates the median of the box; the green triangle indicates the average value of the box. ***, *p*-value < 0.001. UV_irradiation: ultraviolet irradiation condition; ΔDr_ebfC: ΔDr_ebfC mutant strain.

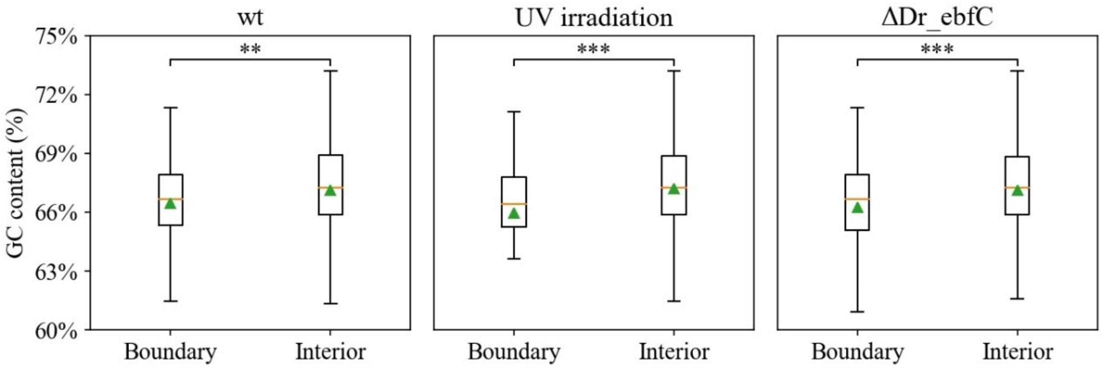


**Figure S3.** Box plots of GC content distributions corresponding different samples. The GC content of CID boundary is significantly lower than that of the CID interior. The orange line indicates the median of the box; the green triangle indicates the average value of the box. **, *p*-value < 0.01; ***, *p*-value < 0.001. wt: wild-type; UV irradiation: ultraviolet irradiation condition; ΔDr_ebfC: ΔDr_ebfC mutant strain.

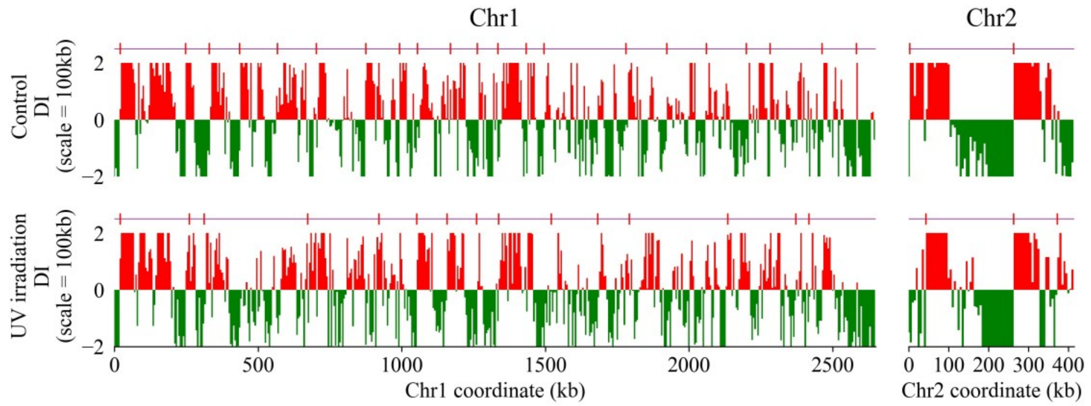


**Figure S4.** Domain boundaries characterized for the control and UV irradiation conditions using a DI analysis performed at a scale of 100 kb. Downstream (red) and upstream (green) biases are indicated. Significant boundaries defining CIDs are annotated with red vertical lines above the panel.

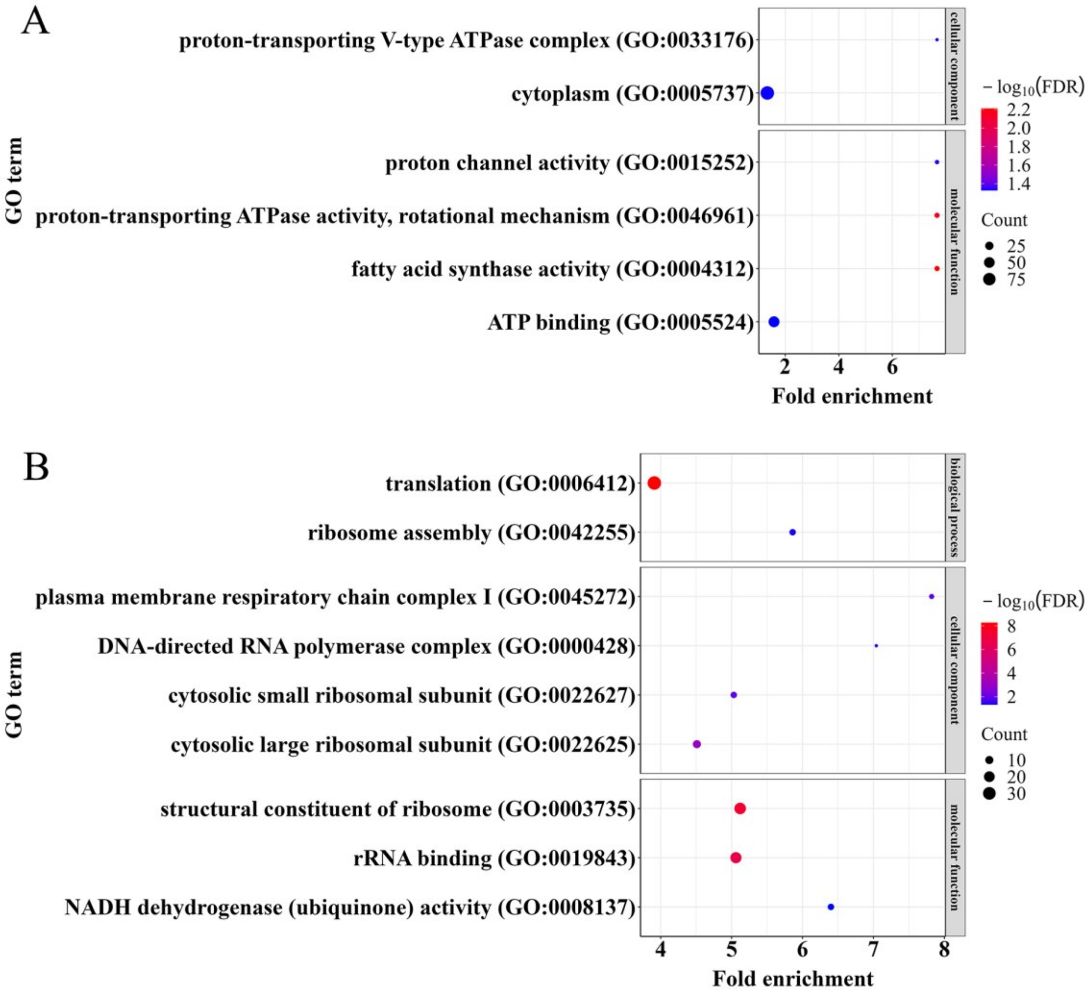


**Figure S5.** Comparison of GO enrichment terms of specific CID boundary genes under two conditions. **(A), (B)** GO enrichment results of the CID boundary genes for the control group and UV irradiation condition, respectively.

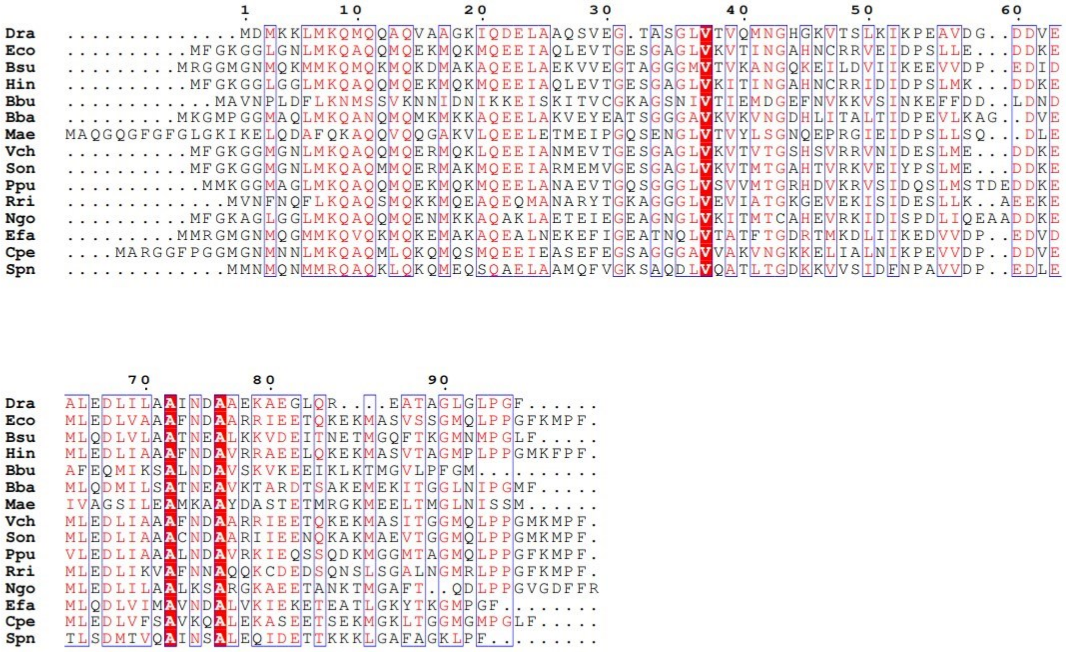

**Figure S6.** Multiple sequence alignment of YbaB/EbfC proteins from different bacterial species using ClustalW software. Identical amino acids are boxed as white characters on a red background, and similar amino acids as red characters on a white background. The selected species are *Deinococcus radiodurans* (Dra), *Escherichia coli* (Eco), *Bacillus subtilis* (Bsu), *Haemophilus influenzae* (Hin), *Borreliella burgdorferi* (Bbu), *Bdellovibrio bacteriovorus* (Bba), *Microcystis aeruginosa* (Mae), *Vibrio cholerae* (Vch), *Shewanella oneidensis* (Son), *Pseudomonas putida* (Ppu), *Rickettsia rickettsiae* (Rri), *Neisseria gonorrhoeae* (Ngo), *Enterococcus faecalis* (Efa), *Clostridium perfringens* (Cpe) and *Streptococcus pneumoniae* (Spn), respectively.

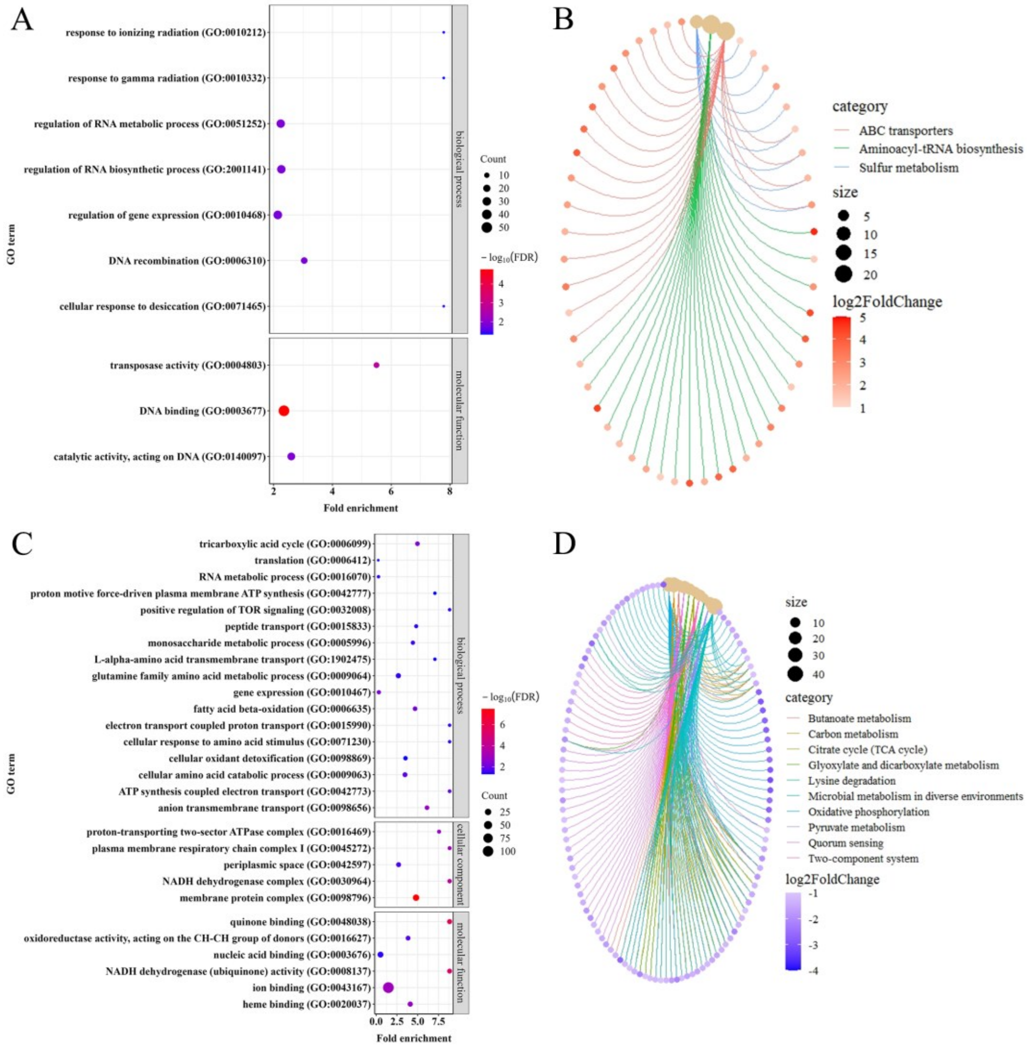


**Figure S7.** Enrichment results of differentially expressed genes (DEGs) in the transcriptional profiles of wild-type and ΔDr_ebfC mutant strains. (A), (B) GO and KEGG enrichment results of the upregulated DEGs, respectively. (C), (D) GO and KEGG enrichment results of the downregulated DEGs, respectively.

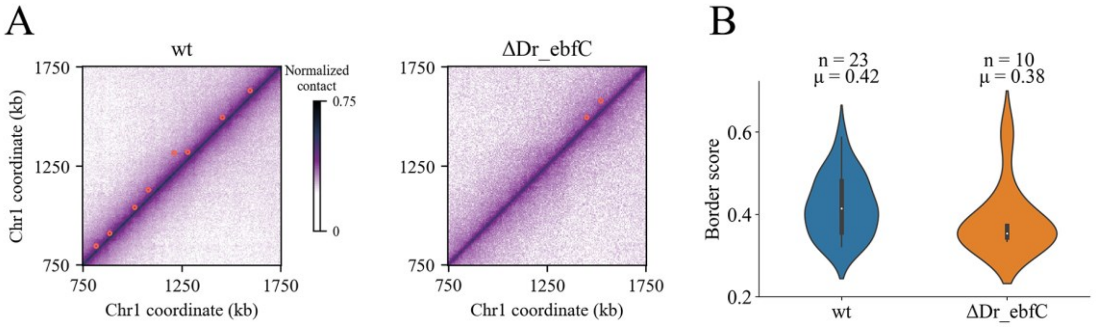


**Figure S8.** Comparison of loop position and strength obtained using Chromosight. (A) Magnification of the 1-Mb region of the Chr1 (coordinates: 750-1750 kb) for wild-type and ΔDr_ebfC mutant cells. The positions of chromosomal loops are indicated in red dots. (B) The number of loops detected and the mean value of loop strength for each cell type (wt and ΔDr_ebfC) are shown on the top.

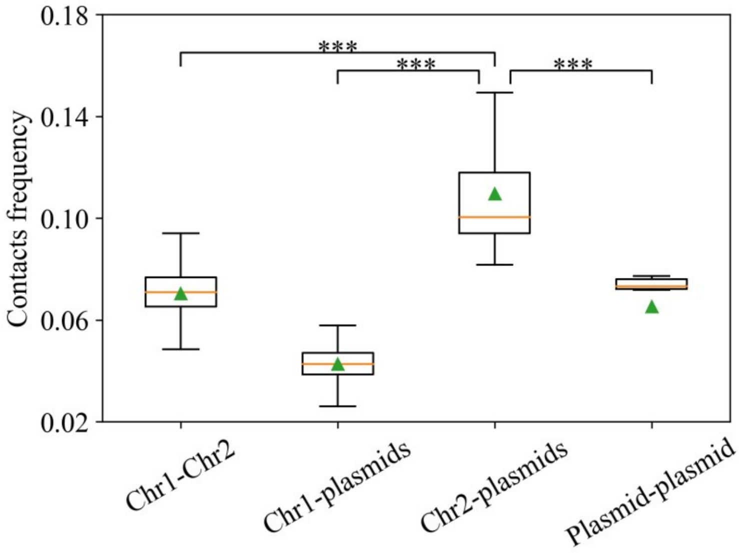


**Figure S9.** Boxplot of contact frequencies for different types of interactions. The orange line indicates the median of the box. The green triangle indicates the average value of the box. ***, $p$-value < 0.001.

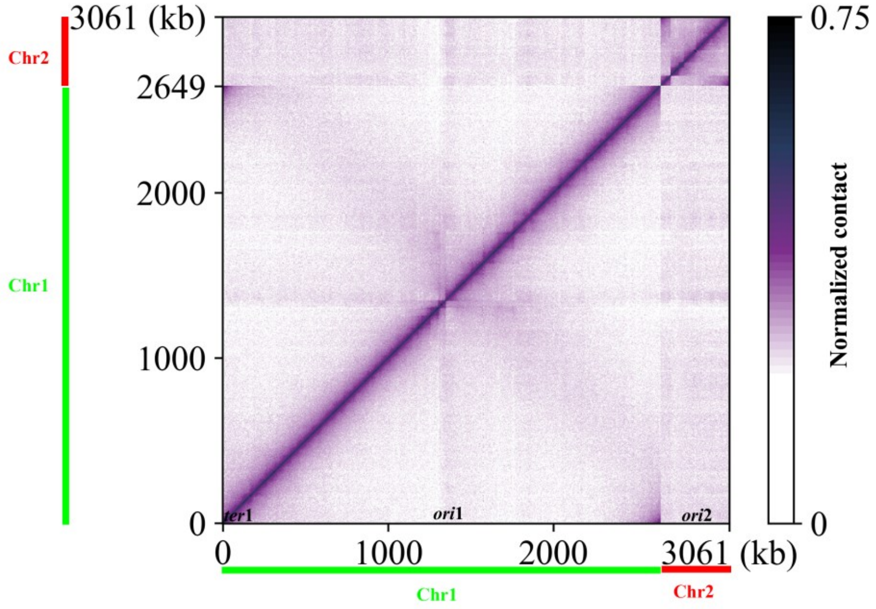


**Figure S10.** Contact map of the two chromosomes of *D. radiodurans*. Chr1 and Chr2 are indicated by green and red bars, respectively. To better visualize contacts in the origin region, the genome coordinates of two chromosomes are rearranged with the origin at the center and the two arms on either side. The positions of the origins (ori1 and ori2) and terminus (ter1) are indicated on the *x* axis.

### References


1. Eugénie N, Zivanovic Y, Lelandais G, Coste G, Bouthier de la Tour C, Bentchikou E, Servant P, Confalonieri F. 2021. Characterization of the radiation desiccation response regulon of the radioresistant bacterium *Deinococcus radiodurans* by integrative genomic analyses. Cells 10:2536.
